# Integrated Proteomics Reveals Brain-Based Cerebrospinal Fluid Biomarkers in Asymptomatic and Symptomatic Alzheimer’s Disease

**DOI:** 10.1101/806752

**Authors:** Lenora Higginbotham, Lingyan Ping, Eric B. Dammer, Duc M. Duong, Maotian Zhou, Marla Gearing, Erik C.B. Johnson, Ihab Hajjar, James J. Lah, Allan I. Levey, Nicholas T. Seyfried

## Abstract

Alzheimer’s disease (AD) features a complex web of pathological processes beyond amyloid accumulation and tau-mediated neuronal death. To meaningfully advance AD therapeutics, there is an urgent need for novel biomarkers that comprehensively reflect these disease mechanisms. Here we applied an integrative proteomics approach to identify cerebrospinal fluid (CSF) biomarkers linked to a diverse set of pathophysiological processes in the diseased brain. Using multiplex proteomics, we identified >3,500 proteins across 40 CSF samples from control and AD patients and >12,000 proteins across 48 postmortem brain tissues from control, asymptomatic AD (AsymAD), AD, and other neurodegenerative cases. Co-expression network analysis of the brain tissues resolved 44 protein modules, nearly half of which significantly correlated with AD neuropathology. Fifteen modules robustly overlapped with proteins quantified in the CSF, including 271 CSF markers highly altered in AD. These 15 overlapping modules were collapsed into five panels of brain-linked fluid markers representing a variety of cortical functions. Neuron-enriched synaptic and metabolic panels demonstrated decreased levels in the AD brain but increased levels in diseased CSF. Conversely, glial-enriched myelination and immunity panels were highly increased in both the brain and CSF. Using high-throughput proteomic analysis, proteins from these panels were validated in an independent CSF cohort of control, AsymAD, and AD samples. Remarkably, several validated markers were significantly altered in AsymAD CSF and appeared to stratify subpopulations within this cohort. Overall, these brain-linked CSF biomarker panels represent a promising step toward a physiologically comprehensive tool that could meaningfully enhance the prognostic and therapeutic management of AD.

## Introduction

Cerebrospinal fluid (CSF) has become one of the most promising sources for accessible biomarkers of neurodegenerative disease. By maintaining direct contact with the brain, CSF has a unique advantage over plasma, saliva, and other fluid sources in its ability to reflect biochemical changes occurring at the core of neuropathology [1]. Alzheimer’s disease (AD), the most common neurodegenerative disorder worldwide, has several well-established CSF markers with direct links to brain-based pathology. These include a) Aβ_1-42_, reflective of cortical amyloid plaque formation; b) Total Tau, a marker of axonal degeneration; and c) Phospho-Tau, representative of pathological tau protein hyperphosphorylation in the AD brain. Together, these “core” CSF markers have significantly advanced AD diagnostics [2, 3]. Yet, despite their diagnostic utility, these three markers reflect only a portion of the complex cellular and molecular dysfunction occurring in the AD brain. A plethora of molecular alterations have now been observed in the cortex of Alzheimer’s patients, extending well beyond amyloid accumulation and tau-mediated neuronal dysfunction [4–6]. Moreover, it is increasingly clear that this complex pathophysiology has important implications for the successful monitoring and treatment of disease. Indeed, the repeated failures of amyloid-targeting therapies have demonstrated that additional pathological pathways are likely critical to the neurodegeneration that leads to clinical decline. Thus, identifying markers of these dysfunctional processes in the earliest asymptomatic stages of disease has become a priority of the AD scientific community [3, 7].

Given this urgent need for markers that more comprehensively reflect AD neuropathology, holistic systems-based approaches have increasingly emerged to address biomarker discovery with the goal of developing marker panels that encompass wider ranges of disease physiology [4, 5]. Network-based proteomics is one such means of mapping the intricate biological systems involved in the pathogenesis of AD and other complex diseases. Our group has previously applied a systems-based proteomic approach to the AD brain and demonstrated across multiple datasets its ability to identify altered networks of protein co-expression linked to specific cell types, organelles, and biological pathways [8–13]. In the current study, we integrated these brain-derived protein systems with analysis of the AD spinal fluid proteome to identify a group of CSF markers reflective of a broad spectrum of molecular alterations in the AD cortex. We ultimately identified and validated five panels of promising CSF targets linked to multiple dysregulated protein systems in the AD brain, including synaptic transmission, vascular biology, myelination, glial-mediated inflammation, and energy metabolism. In addition, by examining these validated targets in the spinal fluid of asymptomatic AD (AsymAD) cases, we were able to identify markers potentially capable of stratifying disease risk among individuals in the earliest stages of illness. Overall, these results represent a critical step toward the translation of systems-based proteomics into accessible biomarker panels that could meaningfully enhance the prognostic and therapeutic management of Alzheimer’s patients.

## Results

### CSF Proteome Reveals Markers Significantly Altered in AD

The main objective of this study was to use an unbiased integrative proteomic approach to identify novel AD fluid biomarkers reflective of the complex systems-based pathology of diseased brain tissue. To achieve this goal, we analyzed a total of four proteomic datasets, including two derived from brain tissue (Brain1 and Brain2) and two from spinal fluid (CSF1 and CSF2). All four cohorts comprised AD dementia samples. Yet, CSF2 and Brain2 were distinguished by their additional inclusion of asymptomatic AD (AsymAD) cases and ultimately allowed us to assess the relevance of our results in this presymptomatic population. **Figure 1** provides an overview of our analytical approach, which comprised a) an initial discovery-driven integrative analysis of brain and spinal fluid proteomes to identify CSF targets of interest in AD dementia (AD) (**Figure 1A**) followed by b) independent validation of these markers in the spinal fluid of both AsymAD and AD subjects (**Figure 1B**). As shown, the integrative portion began with a differential expression analysis of the CSF1 proteome. This proteome was derived from 40 samples comprised of cognitively normal (n=20) and AD (n=20) cases. The clinical diagnoses of AD were supported by cognitive impairment (avg MoCA=13.8), as well as low Aβ_1-42_ and elevated Total Tau and Phospho-Tau ELISA levels in the CSF (**Table 1**). The cognitively unimpaired control cases (avg MoCA=26.7) were selected for normal levels of amyloid and tau in the CSF to verify the absence of preclinical disease pathology.

**Figure 1.**
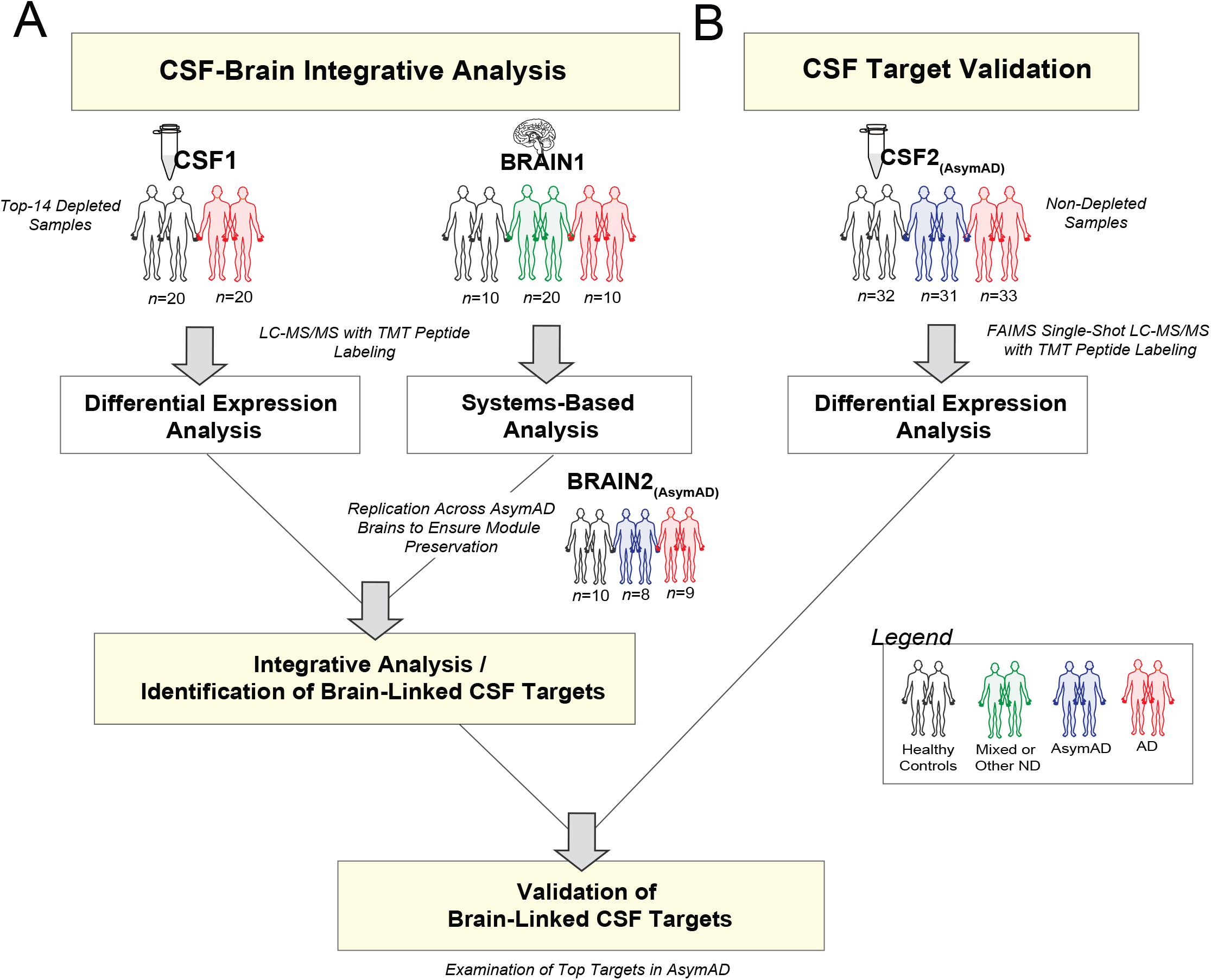
Study Approach. A) Integrative analysis of CSF1 and Brain1 proteomes. The CSF1 proteome comprised 20 healthy control and 20 AD samples, which were each depleted of the fourteen most abundant spinal fluid proteins prior to proteomic analysis. The Brain1 proteome also comprised 40 samples but from three different populations, including 10 control, 10 AD, and 20 other or mixed neurodegenerative cases. Both the CSF1 and Brain1 cohorts were analyzed using isobaric tandem mass tag (TMT) liquid chromatography with tandem mass spectrometry (LC-MS/MS). Differentially expressed proteins (*p*(CT-AD<0.05)) were identified in the CSF1 dataset using statistical t-test analysis, while a systems-based analysis was applied to the Brain1 dataset using weighted protein correlation network analysis (WPCNA). Prior to the integrative analysis of these two datasets, the brain-derived results were examined in a second cohort of post-mortem brain samples (Brain2, *n*=27) that included asymptomatic AD (AsymAD) and module preservation between the two brain datasets was confirmed. The overlap between Brain1 co-expression networks and differentially expressed CSF1 markers was then determined in an integrative analysis and used to identify brain-linked CSF AD targets of interest. These markers were then validated in a separate CSF cohort (CSF2). B) Validation analysis of brain-linked CSF targets. The targets of interest identified in the discovery-driven integrative analysis were examined in the CSF2 cohort, which included 32 control, 31 AsymAD, and 33 AD samples. These cases were analyzed using “single-shot” TMT integrated with high-Field Asymmetric Waveform Ion Mobility Mass Spectrometry (FAIMS), which eliminated the need for sample depletion prior to analysis. Markers of interest were then analyzed in this CSF2 proteome to identify those proteins with reproducible abundance changes, as well as those targets with meaningfully altered levels in preclinical disease. Abbreviations: AD, Alzheimer’s Disease; AsymAD, Asymptomatic Alzheimer’s Disease; ND, Neurodegeneration; LC-MS/MS, Liquid Chromatography with Tandem Mass Spectrometry; TMT, Tandem Mass Tag; High-Field Asymmetric Waveform Ion Mobility Mass Spectrometry, FAIMS.

**Table 1.** Case Characteristics.

|  | CSF1 |  | CSF2 |  |  | BRAIN1 |  |  |  | BRAIN2<sup>a</sup> |
| --- | --- | --- | --- | --- | --- | --- | --- | --- | --- | --- |
|  | CT<br>(n=20) | AD<br>(n=20) | CT<br>(n=32) | AsymAD<br>(n=31) | AD<br>(n=33) | CT<br>(n=10) | PD<br>(n=10) | AD/PD<br>(n=10) | AD<br>(n=10) | AsymAD<br>(n=8) |
| **Age**, mean yrs (SD) | 69.2 (9.0) | 65.6 (11.6) | 62.5 (6.0) | 64.1 (7.9) | 72.6 (7.2) | 63.7 (10.3) | 70.4 (4.5) | 71.6 (8.5) | 65.1 (7.4) | 82.5 (9.7) |
| **Sex**, *n* (%) |  |  |  |  |  |  |  |  |  |  |
| Male | 11 (55.0) | 12 (60.0) | 14 (43.8) | 12 (38.7) | 16 (48.5) | 5 (50.0) | 9 (90.0) | 6 (60.0) | 5 (50.0) | 5 (62.5) |
| Female | 9 (45.0) | 8 (40.0) | 18 (56.2) | 19 (61.3) | 17 (51.5) | 5 (50.0) | 1 (10.0) | 4 (40.0) | 5 (50.0) | 3 (37.5) |
| **Race**, *n* (%) |  |  |  |  |  |  |  |  |  |  |
| White | 12 (60.0) | 13 (65.0) | 17 (53.1) | 22 (71.0) | 26 (78.8) | 7 (70.0) | 10 (100.0) | 8 (80.0) | 10 (100.0) | 8 (100.0) |
| Black | 8 (40.0) | 6 (30.0) | 12 (37.5) | 9 (29.0) | 7 (21.2) | 1 (10.0) | 0 (0.0) | 2 (20.0) | 0 (0.0) | 0 (0.0) |
| Other | 0 (0.0) | 1 (5.0) | 3 (9.4) | 0 (0.0) | 0 (0.0) | 2 (20.0) | 0 (0.0) | 0 (0.0) | 0 (0.0) | 0 (0.0) |
| **PMI**, mean hrs (SD) | -- | -- | -- | -- | -- | 9.3 (7.1) | 6.8 (4.2) | 8.6 (3.9) | 5.4 (1.5) | 17.5 (9.6) |
| **Cognitive Status**,<br>mean (SD)<sup>b</sup> |  |  |  |  |  |  |  |  |  |  |
| MoCA Score | 26.7 (2.2) | 13.8 (7.0) | 26.1 (2.8) | 27.1 (2.2) | 20.0 (3.8) | -- | -- | -- | -- | -- |
| MMSE Score | -- | -- | -- | -- | -- | -- | -- | -- | -- | 28.0 (1.3) |
| **Amyloid & Tau Burden**,<br>mean (SD) |  |  |  |  |  |  |  |  |  |  |
| ELISA<sup>c</sup> |  |  |  |  |  |  |  |  |  |  |
| Aβ<sub>1-42</sub> | 554.3 (90.4) | 223.4 (72.9) | 289.9 (38.5) | 189.6 (45.0) | 144.1 (35.1) | -- | -- | -- | -- | -- |
| Total Tau | 41.0 (17.5) | 143.5 (71.3) | 37.6 (7.6) | 68.1 (30.3) | 111.3 (43.2) | -- | -- | -- | -- | -- |
| Phospho-Tau | 25.2 (8.4) | 57.6 (18.4) | 10.6 (2.9) | 18.5 (11.5) | 35.6 (17.3) | -- | -- | -- | -- | -- |
| CERAD Score | -- | -- | -- | -- | -- | 0.0 (0.0) | 0.3 (0.9) | 3.0 (0.0) | 3.0 (0.0) | 2.8 (0.5) |
| Braak Score | -- | -- | -- | -- | -- | 1.1 (0.7) | 1.6 (0.8) | 5.7 (0.5) | 5.9 (0.3) | 3.1 (1.1) |
| **ApoE Status**, *n* (%)<sup>d</sup> |  |  |  |  |  |  |  |  |  |  |
| 4/4 | 0 (0.0) | 9 (45.0) | 0 (0.0) | 4 (12.9) | 8 (24.2) | 0 (0.0) | 0 (0.0) | 4 (40.0) | 1 (10.0) | 1 (12.5) |
| 3/4 | 4 (20.0) | 4 (20.0) | 6 (18.8) | 10 (32.3) | 17 (51.5) | 2 (20.0) | 0 (0.0) | 4 (40.0) | 6 (60.0) | 2 (25.0) |
| 3/3 | 12 (60.0) | 6 (30.0) | 21 (65.6) | 13 (41.9) | 6 (18.2) | 8 (80.0) | 6 (60.0) | 2 (20.0) | 2 (20.0) | 3 (37.5) |
| 2/4 | 1 (5.0) | 0 (0.0) | 1 (3.1) | 1 (3.2) | 1 (3.0) | 0 (0.0) | 0 (0.0) | 0 (0.0) | 0 (0.0) | 0 (0.0) |
| 2/3 | 3 (15.0) | 0 (0.0) | 2 (6.3) | 2 (6.5) | 1 (3.0) | 0 (0.0) | 4 (40.0) | 0 (0.0) | 1 (10.0) | 1 (12.5) |
| 2/2 | 0 (0.0) | 0 (0.0) | 2 (6.3) | 1 (3.2) | 0 (0.0) | 0 (0.0) | 0 (0.0) | 0 (0.0) | 0 (0.0) | 1 (12.5) |

<sup>a</sup>The Brain2 dataset, which comprised a total 27 tissues, included a re-analysis of 10 control and 9 AD cases from the Brain1 cohort. One of the Brain1 AD cases could not be re-analyzed in Brain2 due to insufficient sample amount. The AsymAD brain samples were unique only to the Brain2 cohort. <sup>b</sup>Two of the AsymAD cases in Brain2 did not have MMSE scores available prior to death but were clinically deemed normal. Neither MOCA nor MMSE scores were available for samples in the Brain1 cohort. <sup>c</sup>The ELISA measurements for CSF1 and CSF2 were obtained on two different assay platforms. CSF1 samples, collected from the Emory Cognitive Neurology Clinic, were sent to Athena Diagnostics and assayed using the INNOTEST® assay platform, while CSF2 samples, collected from research participants of the Emory Goizueta ADRC, Emory Healthy Brain Study, and Emory M<sup>2</sup>OVE-AD study, were assayed using the INNO-BIA AlzBio3Luminex assay. This accounts for the different ranges of Aβ<sub>1-42</sub>, Total Tau, and Phospho-Tau between the two CSF cohorts. <sup>d</sup>Genotype information was missing for one of the AD cases in the CSF1 cohort. Abbreviations: CT, Control; AD, Alzheimer's Disease; AsymAD, Asymptomatic Alzheimer's Disease; PD, Parkinson's Disease; PMI, Postmortem Interval; MoCA, Montreal Cognitive Assessment; MMSE, Mini-Mental State Examination; CERAD, Consortium to Establish a Registry for Alzheimer's Disease; SD, Standard Deviation.

Human spinal fluid is characterized by a dynamic range of protein abundance, in which albumin and other exceedingly high-abundant proteins can prevent the detection of proteins of interest [14]. Thus, to increase the depth of protein discovery, we applied a protocol to each sample designed to deplete fourteen highly abundant spinal fluid proteins, including albumin [14]. Following this depletion, each sample was individually analyzed using quantitative liquid chromatography coupled to tandem mass spectrometry (LC-MS/MS). In total, 39,805 peptides mapping to 3,691 protein groups were identified across the 40 samples. Quantification of proteins was performed via multiplex tandem mass tag (TMT) labeling [9, 15, 16]. To account for missing data, only those proteins quantified in at least 50% of samples were included in subsequent analyses, resulting in the final quantification of 2,875 protein groups. Due to notable differences in total protein abundance levels, one control sample was determined to be an outlier using statistical methods previously described [10, 17] and was not included in subsequent analyses. As part of our analytic pipeline, the abundance values of the remaining 39 samples were adjusted for age, sex, and batch covariance.

Differential expression was then assessed using a statistical t-test analysis, which identified those proteins with significantly altered abundance levels (*p*<0.05) between the control and AD cases (**Table S1**). As demonstrated in the volcano plot of **Figure 2A**, there were a total of 225 proteins with significantly decreased abundance in AD and 303 proteins with significantly increased AD levels. These differentially expressed proteins included several previously identified CSF AD markers, such as microtubule-associated protein tau (MAPT, *p*=3.52E-08), neurofilament light (NEFL, *p*=6.56E-03), growth associated protein 43 (GAP43, *p*=1.46E-05), fatty acid binding protein 3 (FABP3*, p*=2.00E-05), chitinase 3 like 1 (CHI3L1, *p*=4.44E-06), neurogranin (NRGN, *p*=3.43E-04), and VGF nerve growth factor (VGF, *p*=4.83E-03) [2, 18–23]. Yet, we also identified strongly altered novel targets, such as GDP dissociation inhibitor 1 (GDI1, *p*=1.54E-10) and SPARC related modular calcium binding 1 (SMOC1, *p*=6.93E-09). Gene ontology (GO) analysis of the 225 significantly decreased proteins revealed strong biological and functional links to various humoral processes, such as steroid metabolism, blood coagulation, and hormone activity (**Figure 2B, Table S2**). Conversely, the GO terms most highly correlated to the 303 increased proteins indicated strong associations to cell structure and energy metabolism. As expected, MAPT was among the most significantly increased AD proteins and its proteomic levels correlated strongly to independently measured ELISA tau levels (r=0.65, *p*=7.5E-06, **Figure 2C**). Isoform-specific peptides mapping to the C-terminus of Aβ_1-40_ and Aβ_1-42_ do not ionize efficiently following tryptic digestion of amyloid precursor protein (APP) [24, 25]. Therefore, the APP peptides we identified were not used to correlate ELISA Aβ_1-42_ levels.

**Figure 2.**
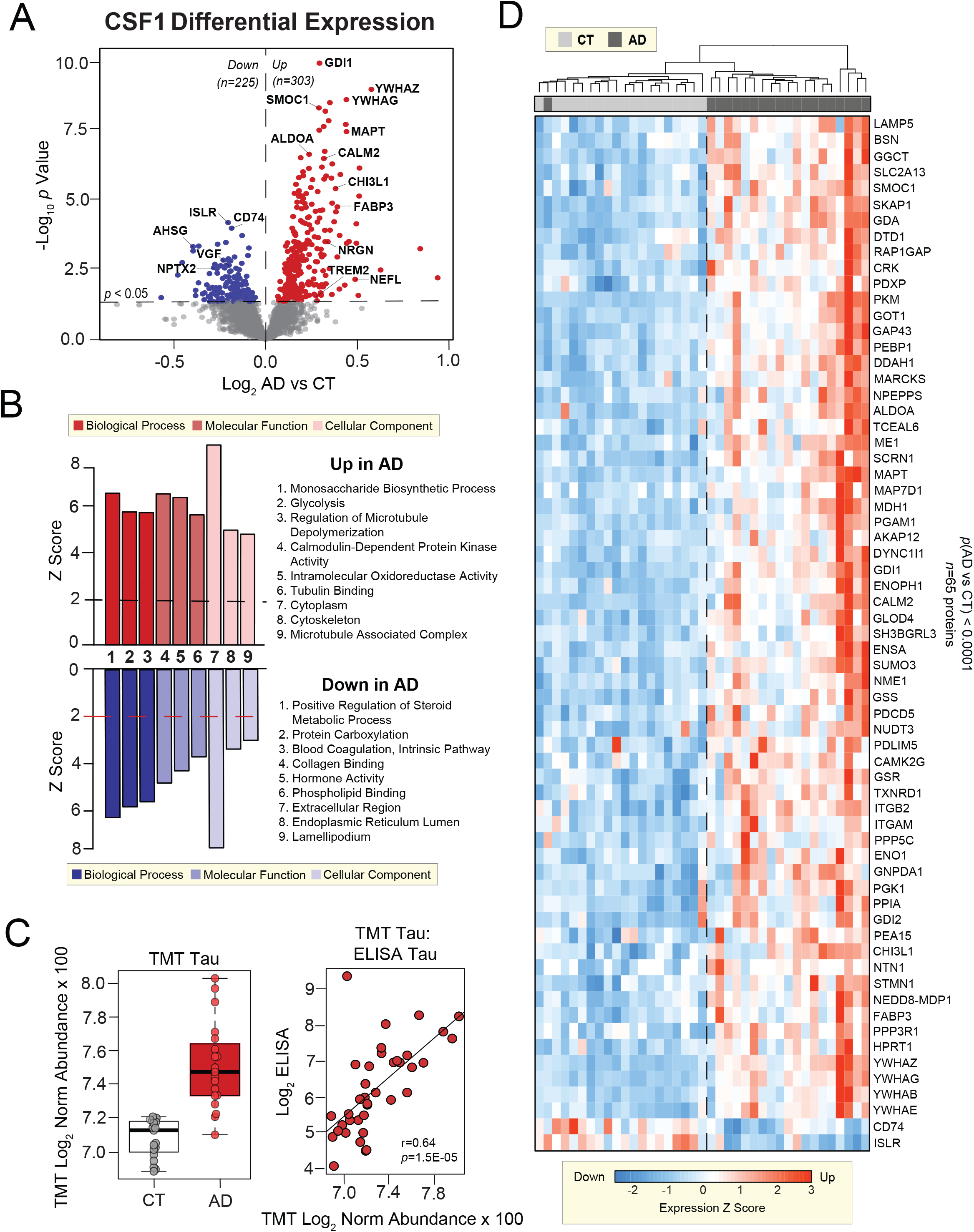
Differential Expression Analysis of the CSF Proteome. A) Volcano plot displaying the log_2_ fold-change (x-axis) against the −log_10_ statistical *p* value for all proteins differentially expressed between control and AD cases of the CSF1 proteome. Those proteins with significantly decreased expression (*p*<0.05) are shown in blue, while the proteins with significantly increased expression (*p*<0.05) are noted in red. Both proteins previously linked to AD (MAPT, CHI3L1, FABP3, NRGN, TREM2, NEFL, VGF, NPTX2) and additional targets (AHSG, ISLR, CD74, GDI1, SMOC1, ALDOA, etc.) are labeled. B) Top gene ontology (GO) terms associated with proteins significantly decreased (blue) and increased (red) in AD. The three GO terms with the highest z-scores in the domains of biological process, molecular function, and cellular component are shown. C) MAPT levels in the CSF1 samples measured by TMT mass spectrometry (left panel) and their correlations to independently measured ELISA Tau levels in the same cases (right panel). The proteomic and ELISA levels correlated strongly across the analyzed samples as assessed by Pearson correlation coefficient and corresponding *p* value (r=0.64, *p*=1.5E-05). Due to missing ELISA data for one AD case, these plots only include values across 38 of the 39 analyzed cases. D) Supervised cluster analysis across the control and AD CSF1 samples using the 65 most significantly altered proteins in the dataset (*p*<0.0001; BH-corrected *p*<0.01). This analysis resulted in the successful differentiation of control and AD cases, except for only one AD case with a control-like marker profile. Abbreviations: TMT, Tandem Mass Tag; Norm, Normalized; BH, Benjamini-Hochberg.

To further analyze differential expression across individual cases, we performed a supervised cluster analysis across the 39 CSF samples using the most significantly altered CSF1 proteins (*p*<0.0001, False Discovery Rate (FDR) *p*<0.01, **Table S1**). As shown in **Figure 2D**, these 65 highly significant proteins were able to correctly cluster cases by disease status except for only one AD case with a control-like marker profile. Of the 65 proteins included in this analysis, 63 were up-regulated in disease. Only the expression of immune-related proteins CD74 and ISLR were consistently down-regulated among AD cases. Overall, the differential expression analyses of the CSF1 proteome not only highlighted hundreds of potential AD fluid markers, but also indicated the most significant proteins among these targets may prove useful for disease classification.

### Systems-Based Analysis of the Brain Proteome Reveals Modules Linked to AD Neuropathology

As outlined in **Figure 1**, we then proceeded with a network-based analysis of the AD dementia brain proteome (Brain1) using an algorithm called weighted protein co-expression network analysis (WPCNA). WPCNA delineates biologically meaningful modules of proteins based on co-expression patterns in large-scale proteomic datasets [10, 17, 26–30]. We have previously applied WPCNA to multiple proteomes of the AD brain and have successfully linked co-expressed protein modules to clinical and pathological phenotypes of disease [8–12]. In the current study, we again applied WPCNA to brain tissue with the goal of identifying co-expression modules that associate with not only AD phenotypes, but also differentially expressed CSF markers. The dorsolateral prefrontal cortex (DLPFC) samples used in this analysis were derived from the Emory Goizueta Alzheimer’s Disease Research Center (ADRC) brain bank and comprised 1) healthy control (*n*=10), 2) AD dementia (*n*=10), and 3) other/mixed neurodegeneration (*n*=20) cases. The cases in this third cohort included subjects with Parkinson’s disease (PD) (*n*=10) and those with mixed AD/PD pathology (*n*=10). The demographics, methods, and basic quantitative details of this brain proteome have been previously described [16]. Overall, the average age and sex of its control and AD cases were well-matched to those of the CSF1 cohort (**Table 1**). Like the CSF1 cases, the 40 brain tissues of the Brain1 cohort, as well as the 27 tissues of the Brain2 cohort, were analyzed using multiplex TMT-based quantification. Collectively, both brain datasets yielded 227,121 unique peptides mapping to 12,943 protein groups across the 67 samples [16]. In our analysis of the Brain1 samples, only those proteins quantified in at least 50% of cases were included in subsequent investigations, resulting in the quantification of 8,817 protein groups across the 40 cases. The abundance levels were then adjusted for the influences of age, sex, and post-mortem interval (PMI).

Prior to systems-level analysis, we first examined the differential expression of our Brain1 proteome by performing a supervised cluster analysis across the 40 samples. Clustering was based on the expression levels of the 165 most significantly altered proteins (*p* value<0.0001) across two Tukey post-ANOVA pairwise comparisons (AD-CT, AD-PD) (**Figure S1A and S1B, Table S3**). As shown, this cluster analysis was able to sharply delineate the 20 cases harboring AD pathology from the control and PD cases. These results indicated that the abundance changes of these significantly altered proteins were largely specific to the AD brain, as opposed to a non-specific result of neurodegeneration. We then applied WPCNA across the 40 tissues. This systems-level analysis identified 44 modules (M) of co-expressed proteins ranked and numbered according to size from largest (M1; *n*=1,821 proteins) to smallest (M44; *n*=34 proteins) (**Table S4**). The algorithm then clustered these modules according to similarities in expression patterns, as represented by the dendrogram of **Figure 3A**. We then analyzed the association of each module to AD neuropathology (i.e. CERAD, Braak) and cell-type specific markers (**Figure 3B**). As previously described [10], cell-type associations were determined based on the overlap of module proteins with those in cell-type specific proteomes derived from isolated neurons, endothelial, and glial cells of the mouse brain. Overall, there were 17 modules that correlated significantly to AD neuropathology (*p*<0.05), many of which were also strongly associated with specific cell types.

**Figure 3.**
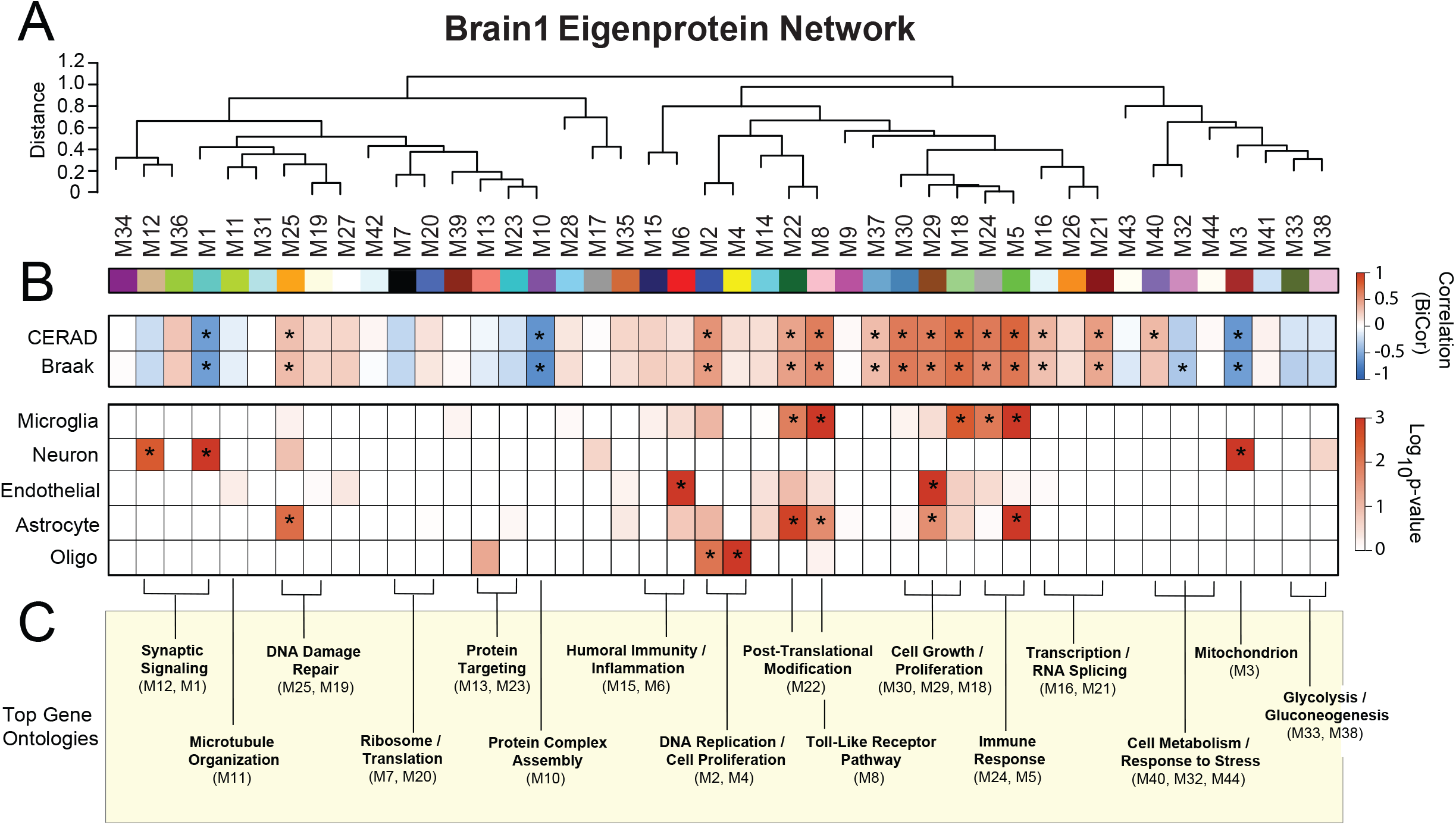
Systems-Based Analysis of the Brain Proteome. A) Weighted protein correlation network analysis (WPCNA) of the Brain1 proteome. This algorithm generated 44 modules (M) of co-expressed proteins. Modules are clustered near each other in the network dendrogram based on their relatedness. B) Biweight midcorrelation (bicor) analysis of module eigenproteins (i.e. the first principle components of module protein expression) with neuropathological hallmarks of AD (top panel). These included CERAD scores, which are reflective of amyloid-β plaque levels in the brain, and Braak scores, which represent brain levels of tau neurofibrillary tangles. In both scales, the higher the score, the greater the amount of pathological burden. Strength of positive (red) or negative (blue) correlation is shown by two-color heatmap with asterisks (*) denoting those statistically significant correlations (*p*<0.05). The cell type associations of each protein module were assessed by module overlap with known microglia, neuron, endothelial, astrocyte, and oligodendrocyte markers. A hypergeometric Fisher exact test (FET) revealed several modules enriched with cell type-specific markers, with the strength of red shading indicating degree of enrichment and statistical significance (*p*<0.05) denoted by asterisks. The FET-derived *p* values were corrected using the Benjamini-Hochberg (BH) method. C) Gene ontology analysis of module proteins. The most strongly associated biological processes are shown for each module or group of related modules. Abbreviations: CERAD, Consortium to Establish a Registry for Alzheimer’s Disease; M, Module; BiCor, Biweight Midcorrelation; Oligo, Oligodendrocyte.

There was notably a cluster of five closely related modules (M30, M29, M18, M24, M5) with strong positive correlations to AD neuropathology. These modules were heavily enriched with microglial and/or astrocytic markers and ontologically linked to cell growth, proliferation, and immunity (**Figure 3C, Table S5**). M8 and M22 were also positively correlated to amyloid and tau burden, enriched with microglia / astrocytes, and linked to the brain’s innate immunity. M8 was highly associated with the toll-like receptor pathway, a signaling cascade that plays a critical role in the innate immune response [31]. Meanwhile, M22 was strongly linked to post-translational modification, a critical regulatory component of innate immunity and inflammation [32]. Finally, M2 was yet another notable module with positive correlations to AD pathology. This oligodendrocyte-enriched module demonstrated ontological links to nucleoside synthesis and DNA replication, suggesting heightened cell proliferation in disease. Overall, these findings supported the increased abundance of modules enriched with glial markers that we have observed in prior AD proteomes [10], potentially reflecting the immune-mediated glial activation widely implicated in AD pathogenesis [33]. Of note, many of these AD-associated glial modules in the current proteome showed lower expression levels in control and PD cases compared to AD, suggesting a disease-specific relationship to AD (**Figure S1C**).

In contrast, there were only four modules (M1, M3, M10, M32) in our Brain1 proteome with significant negative correlations to AD pathology (*p*<0.05), none of which were associated with glial markers (**Figure 3B**). M1 and M3 were instead enriched with neuronal markers. Yet, while M1 was highly associated with synaptic signaling, M3 was strongly linked to mitochondrial ontologies. Though not associated with neuronal markers, M32 mirrored M3 in its ontological links to cellular metabolism. In stark contrast, M10 was highly related to cell growth and the assembly of protein complexes. All four of these modules were increased in controls and PD compared to AD, conferring disease specificity to their pathological associations (**Figure S1C**). Overall, despite the variable characteristics of these four modules, these results aligned well with the decreased abundance of neuronal-linked modules we have previously observed in the AD brain [10]. Furthermore, these findings support the large body of evidence implicating synaptic dysfunction and cellular hypometabolism in AD pathogenesis [34–36], as well as provide a global perspective on their underlying protein alterations in brain. In summary, systems-level analysis of our Brain1 proteome generated modules of protein co-expression with disease-specific alterations in AD consistent with many of our prior findings.

### Analysis of the Asymptomatic AD Brain Proteome Reveals Presymptomatic Network Changes

Alzheimer’s disease is characterized by an early, asymptomatic phase (AsymAD) in which individuals exhibit amyloid accumulation in the absence of clinical cognitive decline [37–42]. This preclinical stage represents a critical window for early detection and intervention. Since we aimed to identify biomarkers relevant to this key stage of disease, we examined whether the modules we derived from our Brain1 proteome were preserved in AsymAD brains. We performed this analysis in 27 DLPFC tissues comprising control (*n*=10), AsymAD (*n*=8), and AD (*n*=9) cases. The control and AD samples were among those included in the analysis of the Brain1 proteome (**Table 1**), while the AsymAD cases were unique to only the Brain2 cohort. These AsymAD cases, also derived from the Goizueta ADRC brain bank, featured abnormally high amyloid levels (avg CERAD = 2.8) despite normal cognition proximate to death (**Table 1**). The Brain2 tissues were analyzed by TMT-MS in a similar manner to the Brain1 cases. However, MS2-based reporter ion quantitation was performed instead of the MS3 quantitation used to analyze the Brain1 proteome. This allowed us to generate deeper coverage in our Brain2 proteome (∼164,000 peptides) compared to the Brain1 dataset (∼95,000 peptides).

Like the analysis of the Brain1 tissues, only those proteins quantified in at least 50% of samples were included in subsequent investigations, resulting in the quantification of 11,244 protein groups. This dataset comprised nearly all the proteins detected in our Brain1 analysis (**Figure S2A**). Of these, 450 proteins demonstrated significantly altered AD abundance levels (*p*(CT-AD)<0.05) in both datasets (**Table S6**). These 450 markers were highly consistent in their direction of change between proteomes (r=0.94; *p*<1.0E-200) (**Figure S2B**). Among increased proteins, those featuring the most concordant changes between datasets were largely members of the glial-enriched M5 and M18 modules of the Brain1 proteome (MDK, COL25A1, MAPT, NTN1, SMOC1, GFAP). Among the decreased proteins, those with the most concordant changes seemed to almost exclusively represent the synapse-associated M1 module (NPTX2, VGF, RPH3A). Module preservation analysis (**Figure S2C**) demonstrated that approximately 80% of protein modules (36/44) in the Brain1 proteome were significantly conserved (*p*<0.05) in the Brain2 dataset, consistent with our previous proteomic network studies [10, 11]. Fourteen of these modules were highly preserved (*p*<0.01) between the two proteomes. The high agreement in differential expression and module composition between the Brain1 and Brain2 proteomes supported the validity of our systems-based approach. Furthermore, the module preservation indicated that the co-expression modules identified in our Brain1 analysis were largely present in AsymAD.

A more detailed analysis of differential expression in the Brain2 dataset highlighted a notable degree of presymptomatic protein alterations, including a total of 151 significantly altered proteins (*p*<0.05) between AsymAD and controls (**Figure S2D**). In agreement with known amyloid burden (i.e. CERAD scores), APP was significantly elevated in both the AsymAD and AD brain. Meanwhile, MAPT was significantly altered only in AD. Glial-enriched modules (M5, M18) were highly reflected among increased AsymAD proteins, while those proteins decreased in AsymAD tended to be members of the neuronal-linked M1 module. As shown, many of these presymptomatic markers demonstrated even greater changes in symptomatic disease. Among such markers was SMOC1, a glial protein linked to brain tumors and eye and limb development, but whose role in neurodegeneration remains largely undefined [43, 44]. In contrast, the synaptic protein neuropentraxin 2 (NPTX2) was significantly decreased in the AsymAD brain. NPTX2 has been previously linked to neurodegeneration and has a well-established role in mediating the excitatory synapse [45, 46]. Overall, these results indicated presymptomatic changes in not only individual markers but also entire modules of co-expressed proteins.

### Integrative Analysis Identifies Panels of Brain-Linked CSF Biomarkers of AD

After separately analyzing the CSF and brain proteomes of AD dementia and ensuring the protein co-expression modules identified in the AD brains were also preserved in AsymAD, we then applied an integrative analysis to CSF1 and Brain1 to identify fluid markers linked to brain network physiology. Integral to this analysis was defining the overlap between the brain and CSF proteomes. While it is well accepted that the CSF mirrors neurochemical changes in the AD brain [1], the precise degree of overlap between the AD brain and CSF is unclear. By comparing the number of shared gene products detected among our two proteomes, we found that nearly 70% (*n*=1,936) of proteins identified in the CSF1 proteome were also quantified in the brain (**Figure 4A**). The bulk of these overlapping proteins (*n*=1,721) were assigned to one of the 44 co-expression modules derived from the Brain1 dataset (**Figure 4B**). As expected, the six largest brain modules (M1-M6) demonstrated the greatest amount of CSF overlap. However, there were smaller brain modules (e.g. M15, M29) that achieved a surprisingly high degree of overlap, greater than brain modules twice their size. This prompted us to take a more detailed, statistically-driven approach to calculating overlap between our Brain1 and CSF1 datasets.

**Figure 4.**
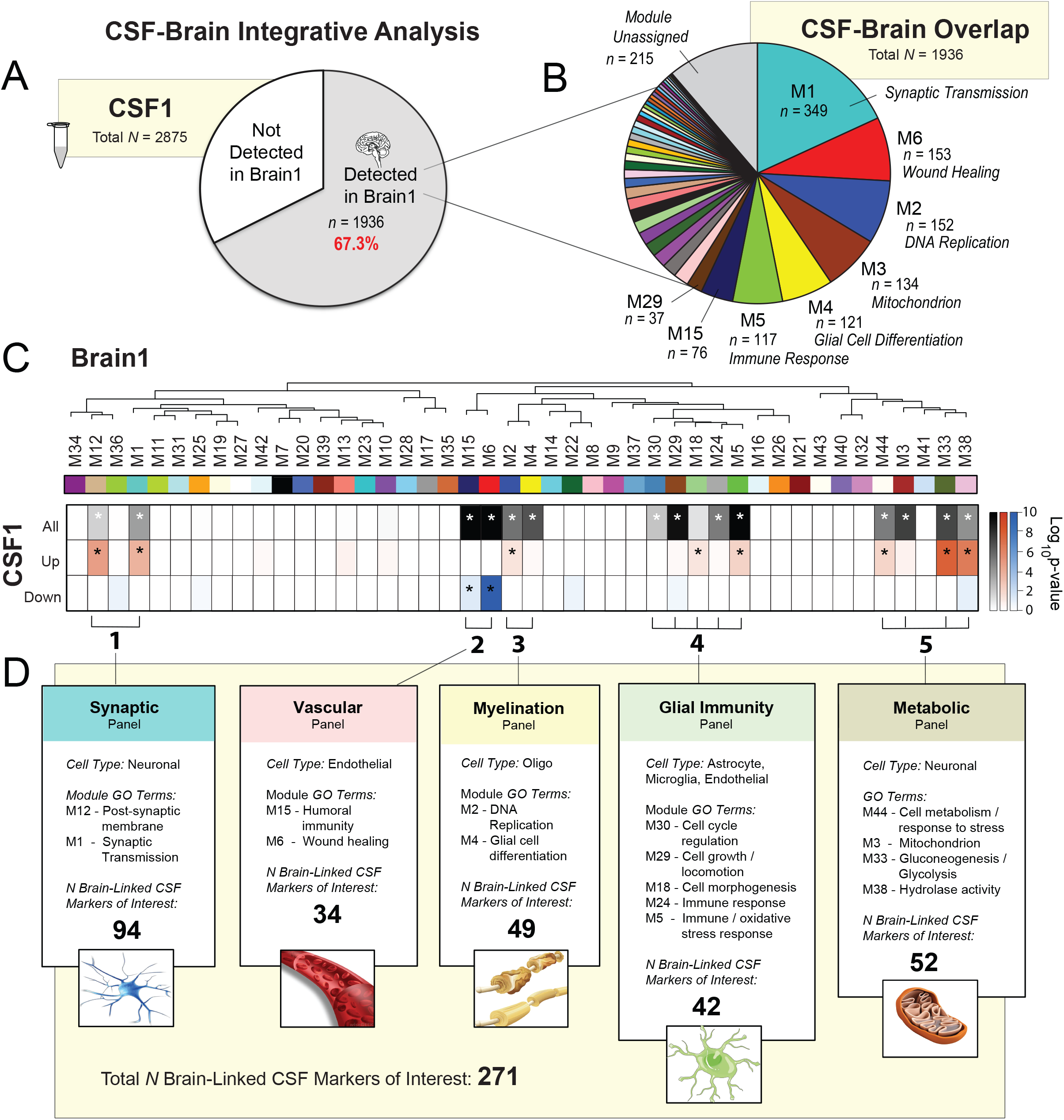
Integrative Analysis of CSF and Brain Proteomes Yields Panels of Brain-Linked CSF AD Biomarkers. A-B) Overlap of proteins detected in the Brain1 and CSF1 datasets. Nearly 70% of proteins quantified in the CSF1 proteome (*n*=1936) were also quantified in the Brain1 proteome. Most of these overlapping proteins were linked to one of the 44 co-expression modules (M) of the Brain1 systems-based network. Modules with the highest numbers of overlapping proteins are labeled. C) Statistical overlap of CSF1 proteins and modules of the Brain1 proteome. Each line of the heat map represents a separate overlap analysis by hypergeometric Fisher’s exact test (FET). The top line depicts the overlap of brain module proteins with the entire CSF1 proteome. This analysis identified 14 modules with significant levels (*p*<0.05) of overlap with quantified CSF1 proteins, denoted with asterisks and gray / black shading. There was also one module (M18) that closely approached significance (*p*=0.06). The second line depicts the overlap of module proteins with the 303 significantly up-regulated (*p*(CT-AD)<0.05) proteins of the CSF1 proteome. Red shading and asterisks highlight the 8 modules (M12, M1, M2, M18, 5, M44, M33, M38) that significantly overlapped (*p*<0.05) with these increased CSF markers. The third line demonstrates brain module overlap with the 225 significantly down-regulated (*p*(CT-AD)<0.05) CSF1 proteins. As denoted by the blue shading and asterisks, only 2 modules (M15, M6) displayed significant overlap (*p*<0.05) with these decreased CSF markers. The FET-derived *p* values were corrected using the Benjamini-Hochberg (BH) method. D) Collapsed module panels based on cell type associations and related gene ontology (GO) terms. Altogether, these panels comprised a total of 271 overlapping proteins with meaningful differential expression in the CSF1 proteome (i.e. brain-linked CSF markers of interest). Abbreviations: M, Module; GO, Gene Ontology; Oligo, Oligodendrocyte.

Using a one-tailed Fisher Exact Test (FET), we assessed the significance of protein overlap between the CSF proteome and individual brain modules. This analysis revealed a total of 14 brain modules with statistically significant overlap in the CSF1 dataset (FDR *p*<0.05), as well as one additional module (M18) whose extent of overlap approached significance (FDR *p*=0.06) (**Figure 4C; top row**). We were also interested in modules that overlapped strongly with differentially expressed CSF proteins. Therefore, we applied two additional FET analyses to determine those brain modules with meaningful overlap among 1) CSF1 proteins significantly increased in AD and 2) CSF1 proteins significantly decreased in AD (*p*[CT-AD] < 0.05). As shown in the middle and bottom rows of **Figure 4C**, these additional analyses revealed 8 of the 44 brain modules significantly overlapped with proteins increased in AD CSF (M12, M1, M2, M18, M5, M44, M33, M38), while only two modules (M6, M15) demonstrated meaningful overlap with proteins decreased in AD CSF. As expected, all 10 of these modules were among the 15 modules with the highest degree of overlap with the total CSF proteome. We therefore hypothesized that the 15 modules with high overlap in the CSF were collectively high-yield sources of brain-linked spinal fluid biomarkers of AD.

We collapsed these 15 overlapping modules of interest into five large protein panels based on their adjacent clustering in the WPCNA dendrogram and associations with cell types and gene ontologies. (**Figure 4D**). The first panel comprised modules strongly enriched with neuronal markers and synapse-associated proteins (M1, M12). This *synaptic* panel contained a total of 94 proteins with significantly altered levels in our CSF1 proteome, making it the largest source of brain-linked CSF markers among our five panels. The second panel (M6, M15) demonstrated strong links to endothelial cell markers and vascular ontologies, such as “wound healing” (M6) and “regulation of humoral immune response” (M15). M15 was also highly associated with lipoprotein metabolism, a process intimately associated with the endothelium [47]. This *vascular* panel harbored 34 proteins brain-linked fluid markers. The third panel comprised modules (M2, M4) significantly linked to oligodendrocyte markers and cellular proliferation. For example, the top ontological terms for M2 included “positive regulation of DNA replication” and “purine biosynthetic process”. Meanwhile, those of M4 included “glial cell differentiation” and “chromosome segregation”. Defined by its strong links to proliferating oligodendrocytes, this *myelination* panel harbored 49 brain-linked fluid markers.

The fourth panel comprised the largest number of modules (M30, M29, M18, M24, M5), nearly all of which were significantly enriched with microglia and astrocyte markers. Like the myelination panel, this fourth panel also contained modules with strong associations to cell proliferation (M30, M29, M18). Yet, its remaining modules were highly associated with immunological terms, such as “immune effector process” (M5) and “regulation of immune response” (M24). This *glial immunity* panel contained 42 brain-linked fluid markers among its overlapping proteins. Finally, the last panel included 52 brain-linked markers over four modules (M44, M3, M33, M38), all of which were ontologically linked to energy storage and metabolism. The largest of these modules (M3) was strongly associated with mitochondria and enriched with neuronal-specific markers. M38, one of the smaller module members of this *metabolic* panel, also demonstrated modest neuronal specificity. Overall, these five panels reflected a wide range of cell types and functions in the AD cortex and collectively harbored 271 brain-linked fluid markers of interest (**Table S7**).

### Synaptic, Vascular, and Metabolic Panels Demonstrate Divergent Expression Trends in the Brain and CSF

The biological themes highlighted by our five panels, from synaptic signaling to energy metabolism, have all been implicated in the pathogenesis of AD [23, 48, 49]. Accordingly, all 15 modules comprising these panels correlated to AD pathology in our Brain1 proteome (**Figure 3B**). Most notable were the highly positive pathological correlations among our glial modules and the strongly negative pathological correlations of our largest neuronal modules (M1, M3). The differential expression analysis of our Brain2 proteome (**Figure S2D**) also highlighted M5 and M18-derived glial proteins among those most increased and M1-associated synaptic proteins among those most decreased in both asymptomatic and symptomatic AD. These observations indicated that the 271 CSF markers we had identified among the five panels were indeed linked to critical disease processes in the AD cortex, including those occurring in early presymptomatic stages.

In order to better resolve the direction of change of panel proteins in the brain and spinal fluid, we plotted the following for each of the 15 overlapping modules: 1) module abundance levels in the Brain1 dataset and 2) the differential expression of module proteins in the CSF1 dataset (**Figure 5**). Module abundance values in the brain were determined using the WPCNA eigenprotein method as previously described [10]. Volcano plots were used to depict the differential expression (CT-AD) of module proteins in the CSF. Interestingly, these plots revealed that three of the five panels demonstrated divergent expression trends in brain and spinal fluid. Both modules of the synaptic panel (M1, M12) demonstrated decreased abundance levels in the AD brain but overlapped significantly with proteins *increased* in AD spinal fluid (**Figure 5A**). The neuronal-associated modules comprising the metabolic panel (M3, M38) demonstrated similarly discordant brain and CSF expression patterns (**Figure 5E**). Finally, the vascular panel also displayed divergent expression trends, though its modules (M6, M15) were modestly increased in the AD brain and starkly *decreased* in diseased CSF (**Figure 5B**). The two remaining panels comprised large glial networks whose proteins were concordantly up-regulated in both compartments (**Figure 5C and 5D**).

**Figure 5.**
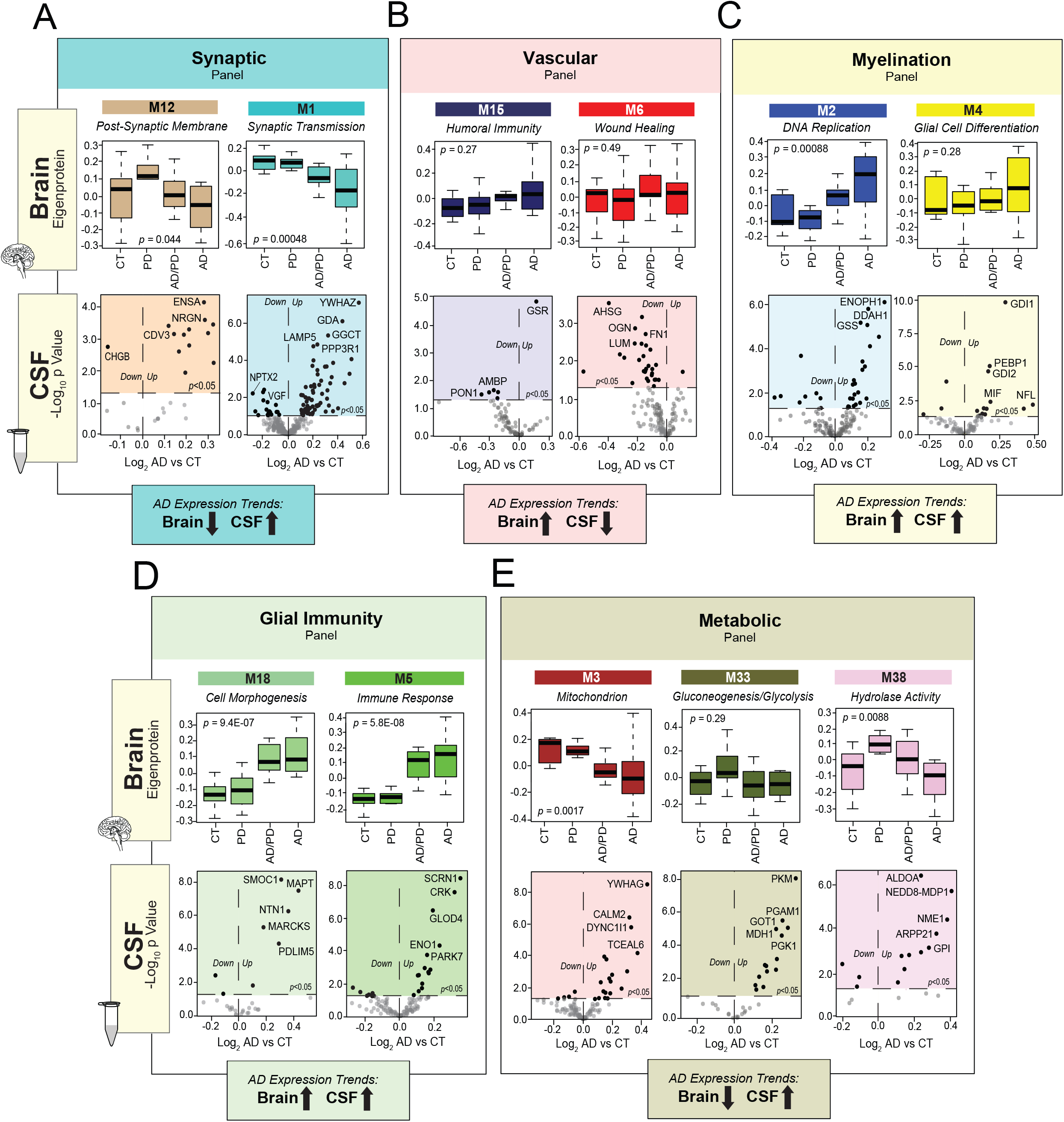
Brain-Linked CSF Marker Panels Demonstrate Divergent Expression Levels in Brain and CSF. A-E) Protein expression trends of the overlapping modules comprising the five marker panels. Module (M) eigenproteins depict expression profiles in the Brain1 proteome, while volcano plots demonstrate trends in the differential expression (log_2_ AD vs CT) of module proteins in the CSF1 proteome. Three panels demonstrated divergent expression trends in the brain and CSF, including the synaptic (A), vascular (B), and metabolic (E) panels. As shown, modules of the synaptic and metabolic panels demonstrated decreased expression in the AD brain but overlapped significantly with spinal fluid proteins up-regulated in AD. The vascular panel featured modules modestly decreased in the AD brain but overlapped significantly with CSF proteins down-regulated in AD. In contrast, the glial-associated panels (C-D) featured modules with concordant up-regulation in the brain and CSF. ANOVA with Tukey post-hoc correction was used to assess the statistical significance of module eigenprotein abundance changes across the four groups of the Brain1 cohort. Abbreviations: CT, Control; PD, Parkinson’s Disease; AD, Alzheimer’s Disease.

It is important to note that these trends were not universal for all markers within these panels. For instance, the synaptic panel included several proteins significantly decreased in the AD brain and CSF (**Figure 5A**). Among these down-regulated CSF markers were NPTX2 and VGF of M1, as well as chromogranin B (CHGB) of M12. Yet, despite these few exceptions, most of our synaptic markers were elevated in AD spinal fluid. Overall, these analyses were able to distinguish statistically meaningful trends in both the brain and spinal fluid levels for each of our five panels. Remarkably, these trends highlighted complex and often divergent relationships between brain and CSF protein expression in AD.

### CSF Biomarkers of Brain-Linked Panels Validate in a Replication Cohort

Our integrative proteomic analysis had identified 271 promising CSF protein markers with links to the AD brain. As a next step, we aimed to narrow this group of proteins to the most informative markers for our final target panels. An ideal AD biomarker is not only able to detect a fundamental feature of disease neuropathology, but also reliable, easy to perform, and capable of recognizing AD early in its course [50]. Indeed, the now widespread recognition that AD neurodegeneration begins years prior to the onset of cognitive symptoms and the growing push to define this disorder principally by physiological markers of neuropathology and neuronal injury has created an especially high demand for preclinical biomarkers [3, 38]. Therefore, we structured our validation analysis to identify which of the 271 targets shared these ideal characteristics. Our first strategy was to include cases at earlier stages of disease to examine the behavior of our markers before the onset of severely devastating cognitive decline. As shown in **Figure 1**, the 96 cases of this CSF2 validation cohort comprised a group of individuals with AsymAD. These subjects were defined by their normal-range cognitive performance (avg MoCA=27.1, Clinical Dementia Rating (CDR)=0) in the setting of pathological levels of core AD biomarkers (**Table 1**). The symptomatic cases of the CSF2 cohort included an earlier spectrum of AD severity, ranging from mild to moderately severe cases. This yielded a notable difference between the average MoCA score of AD cases in CSF2 (20.7) and that of the advanced AD cases included in the CSF1 analysis (13.8).

In addition to early AD biomarkers, we were also interested in protein targets that could be quantified using more efficient, high-throughput mass spectrometry methods that would be necessary in most clinical settings. Therefore, following TMT labeling, we analyzed our CSF2 samples by “single-shot” LC-MS/MS integrated with high-Field Asymmetric Waveform Ion Mobility Mass Spectrometry (FAIMS). When combined with synchronous precursor selection MS3-based quantitation (SPS-MS3), this FAIMS-based strategy is especially useful for enhancing accuracy of protein quantification in samples with high dynamic ranges of protein abundance, such as albumin- and immunoglobulin-rich spinal fluid [51]. Its ion separation step helps to limit the signal interference of albumin and other overwhelmingly abundant proteins, allowing for enhanced quantification of detected low-abundance proteins. This enabled us to circumvent the need for albumin depletion. The application of this more streamlined proteomic approach to the CSF2 cohort resulted in a dataset comprised of 6,487 peptides mapping to 1,183 protein groups across the 96 cases. As in the CSF1 analysis, only those proteins quantified in at least 50% of samples were included in subsequent analyses. This resulted in the final quantification of 792 protein groups. Nearly 95% of the proteins quantified in CSF2 were also identified in the CSF1 proteome. Since we were specifically interested in validating the 271 markers from our integrative analysis, we examined how many of these targets were detected in the CSF2 proteome. We found that 100 of these 271 brain-linked fluid markers were detected in the CSF using both proteomic methodologies. **Figure 6A** demonstrates the differential expression of these 100 overlapping markers across the control and AD individuals from the CSF2 samples (**Table S8**). As illustrated, many of these targets were highly altered in the AD CSF. Those proteins most significantly increased in AD were members of the synaptic (BASP1, HPRT1, YWHAZ) and metabolic (ALDOA, PGK1, GOT1) panels. SMOC1 of the glial immunity panel was also strongly elevated, as well as SOD1 and SPP1 of the myelination panel. Meanwhile, those proteins most decreased in our CSF2 cohort comprised only members of the vascular panel (PON1, AMBP, DCN). Altogether, the 100 overlapping targets demonstrated strong correlations between the two CSF datasets in degree and direction of change in AD (*r*=0.5; *p*=2.3E-05). Indeed, we found that the majority of these 100 proteins (*n*=70) maintained the same directionality of change between the two proteomes. Accordingly, an analysis of marker directionality considering only these 70 proteins revealed even stronger correlations between the two datasets (*r*=0.8; *p*=4.6E-17) (**Figure 6B**). We found these significant correlations particularly notable given the differences in sample preparation, mass spectrometry approaches, and average disease severity between the two AD groups.

**Figure 6.**
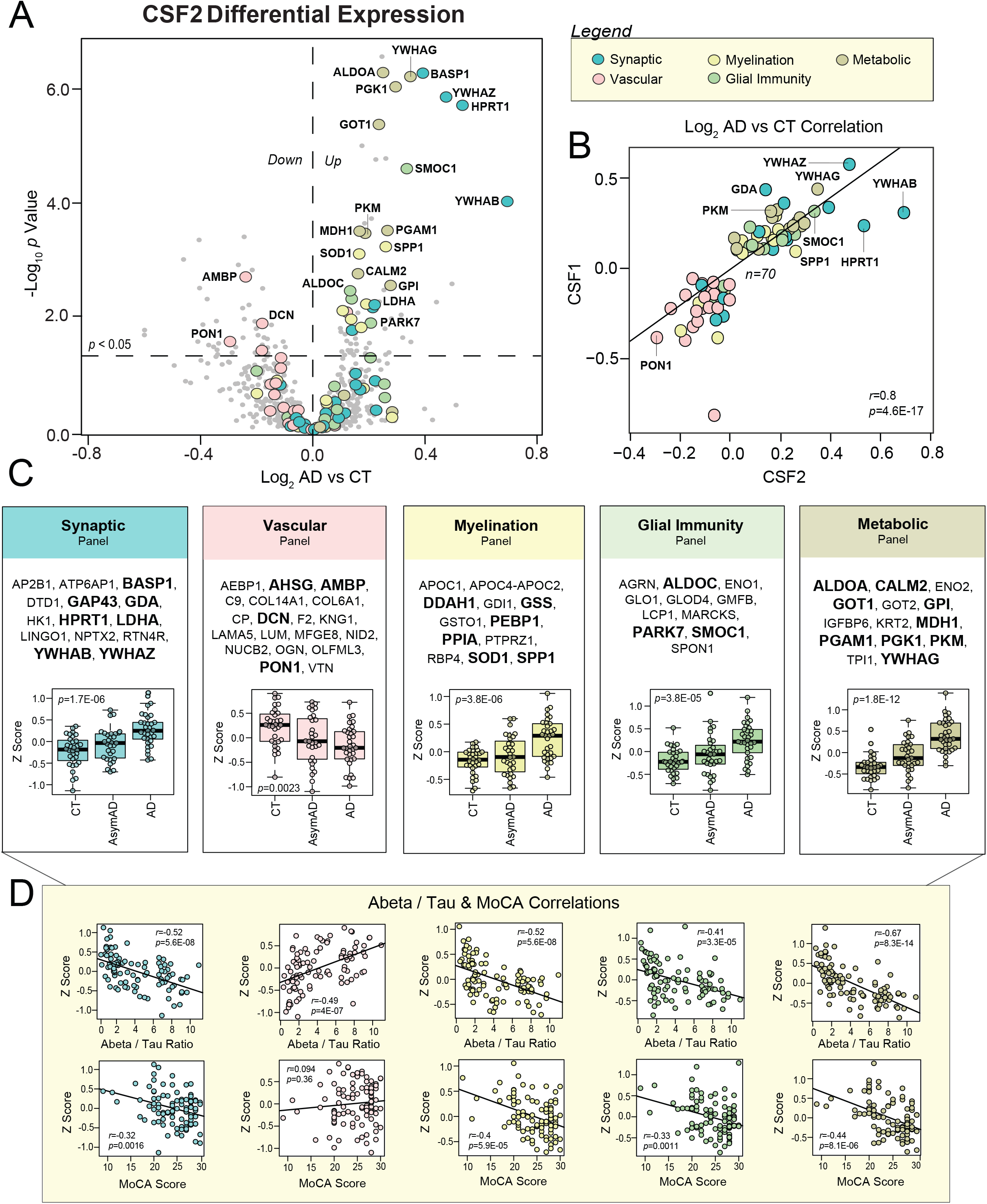
CSF2 Analysis Validates Proteins Across the Five Biomarker Panels. A) Volcano plot displaying the log_2_ fold-change (x-axis) against the −log_10_ statistical *p* value for all proteins differentially expressed between control and AD cases of the CSF2 proteome. Overall, 100 of the 271 panel markers of interest were detected in this dataset. These 100 proteins are represented by the large colored datapoints. Each of the colors reflects a different panel; see figure legend. Of these 100 proteins, 70 demonstrated concordant directionality of change between the two CSF datasets and were deemed validated. B) Correlation analysis of CSF1 and CSF2 differential expression (log_2_ AD vs CT) of the 70 validated protein targets. Pearson correlation coefficient and *p* value were used to assess the degree of correlation. C) Validated protein targets (*n*=70) organized by panel membership. Bolded markers (*n*=29) are those that demonstrated statistically significant AD abundance changes (*p*(CT-AD)<0.05) in both CSF datasets. Panel expression levels were calculated (z-score) across all 96 samples of the CSF2 cohort. ANOVA with Tukey post-hoc correction was used to assess the statistical significance of panel abundance changes across control, AsymAD, and AD groups. D) Panel correlation analyses of protein abundance levels to ELISA Abeta_1-42_/Tau ratios and MoCA scores across the 96 CSF2 cases. Degree of correlation was assessed by Pearson correlation coefficients and corresponding *p* values. Abbreviations: CT, Control; AsymAD, Asymptomatic AD; AD, Alzheimer’s Disease; Abeta, Abeta_1-42_; MoCA, Montreal Cognitive Assessment.

The 70 validated brain-linked CSF markers are highlighted in **Figure 6C**. As shown, all five of our brain network-based panels were represented across these 70 markers. The vascular panel contributed the most proteins (*n*=19) to this validated group, followed by the synaptic panel (*n*=14). Of these 70 proteins, we identified 29 markers with statistically significant AD abundance changes in both datasets. These strongly validated markers are bolded in **Figure 6C**. Among these validated targets were proteins with known connections to AD such as osteopontin (SPP1), a proinflammatory cytokine that has been linked to AD across several studies [52–54]. GAP43 of the validated synaptic panel has also been linked to neurodegeneration and its critical role in synaptic stability has been well-characterized [23, 55]. We also validated many novel markers, such as glial-associated SMOC1 and the synapse-bound BASP1, both of which have very limited prior connections to AD. Interestingly, also among our most strongly validated proteins were markers associated with other neurodegenerative diseases, such as the ALS-linked SOD1 and PD-associated PARK7. On the other hand, we had difficulty detecting filamentous proteins, such as MAPT and NEFL, using our high-throughput validation method and therefore neither of these AD-associated markers were included in our final panels. We were also unable to validate the calmodulin-binding post-synaptic protein, neurogranin (NRGN), which several studies have found elevated in the spinal fluid of AD subjects [20, 21, 36, 56]. NRGN was strongly up-regulated in our CSF1 proteome (**Figure 5A**), but not detectable using the streamlined single-shot proteomics of CSF2. These results align with other inconsistencies in the biofluid detection and measurement of NRGN, possibly stemming from its fragmentation into a series of C-terminal peptides prior to its release into the CSF [56].

While our validated protein biomarkers ultimately reflected a wide array of brain-derived protein systems, a common theme of dysregulated redox potential and energy metabolism seemed to emerge across all five marker panels. In addition to proteins involved in signal transmission (NPTX2, YWHAZ, BASP1), our validated synaptic panel also comprised several targets more classically known for roles in metabolic processes, such as the glycolytic enzyme LDHA and the purine salvaging enzyme HPRT1. This observation likely reflects the close relationship between energy production and synaptic signaling at the neuronal membrane [57, 58]. The validated proteins among our vascular and glial panels also included various markers responsive to oxidative stress. This included SOD1, glutathione synthetase (GSS), and dimethylarginine dimethylaminohydrolase 1 (DDAH1) of the validated myelination panel, which all function to clear damaging free radicals [59–62]. Likewise, the validated glial immunity panel included PARK7 or DJ-1, yet another mediator of oxidative stress in the brain [63, 64]. Finally, highly validated markers of the vascular panel included paroxonase 1 (PON1), a lipoprotein-binding enzyme responsible for reducing oxidative stress levels in the circulation [65, 66]. These results suggested a common functional thread among our most strongly validated markers despite their varied cell type and ontological associations.

The expression levels of our validated panels were calculated (z-score) across all 96 samples of the CSF2 cohort (**Figure 6C**), demonstrating AD abundance trends consistent with those observed in our initial pre-validation panels (**Figure 5**). These expression levels also indicated that our validated markers demonstrated notable changes in presymptomatic disease. Accordingly, all five panels demonstrated strong correlations to the Aβ_1-42_ / Tau ratios of our samples, especially the metabolic panel (*r*=0.67; *p*=8.3E-14) (**Figure 6D**). As expected, the vascular panel was the only one that demonstrated negative correlations to Aβ_1-42_ / Tau ratio (*r*=0.52; *p*=5.6E-08). In contrast, the panel correlations to MoCA were notably weaker, though still statistically significant. Overall, these findings suggested we had validated panels that not only distinguished healthy controls from AD dementia, but also reflected physiological changes occurring in the preclinical stages of disease, prior to the onset of notable cognitive decline.

### Brain-Linked CSF Biomarkers Reveal Heterogeneity Among Asymptomatic AD Cases

It is well-known that individuals with AsymAD are not a homogeneous population and that an interplay of heightened risk and resilience contributes to variability in the subsequent progression of disease [67]. While used to identify asymptomatic AD cases, the levels of core CSF biomarkers (Aβ_1-42_, Total Tau, and Phospho-Tau) have still not demonstrated the ability to reliably predict which individuals will progress to dementia [2, 22]. Therefore, novel biomarkers are also needed to better define and stratify this pre-symptomatic stage of disease. For these reasons, AsymAD samples (*n*=31) were included in our CSF2 cohort. As shown in **Figure 7A**, the 31 AsymAD cases of the CSF2 dataset had Aβ_1-42_, Total Tau, and Phospho-Tau levels that significantly differed from those of both controls and AD patients. In contrast, there was no statistical difference between the MoCA scores of control and AsymAD cases. All individuals with AsymAD also had a CDR of 0, indicating no evidence of decline in everyday cognitive or functional performance.

**Figure 7.**
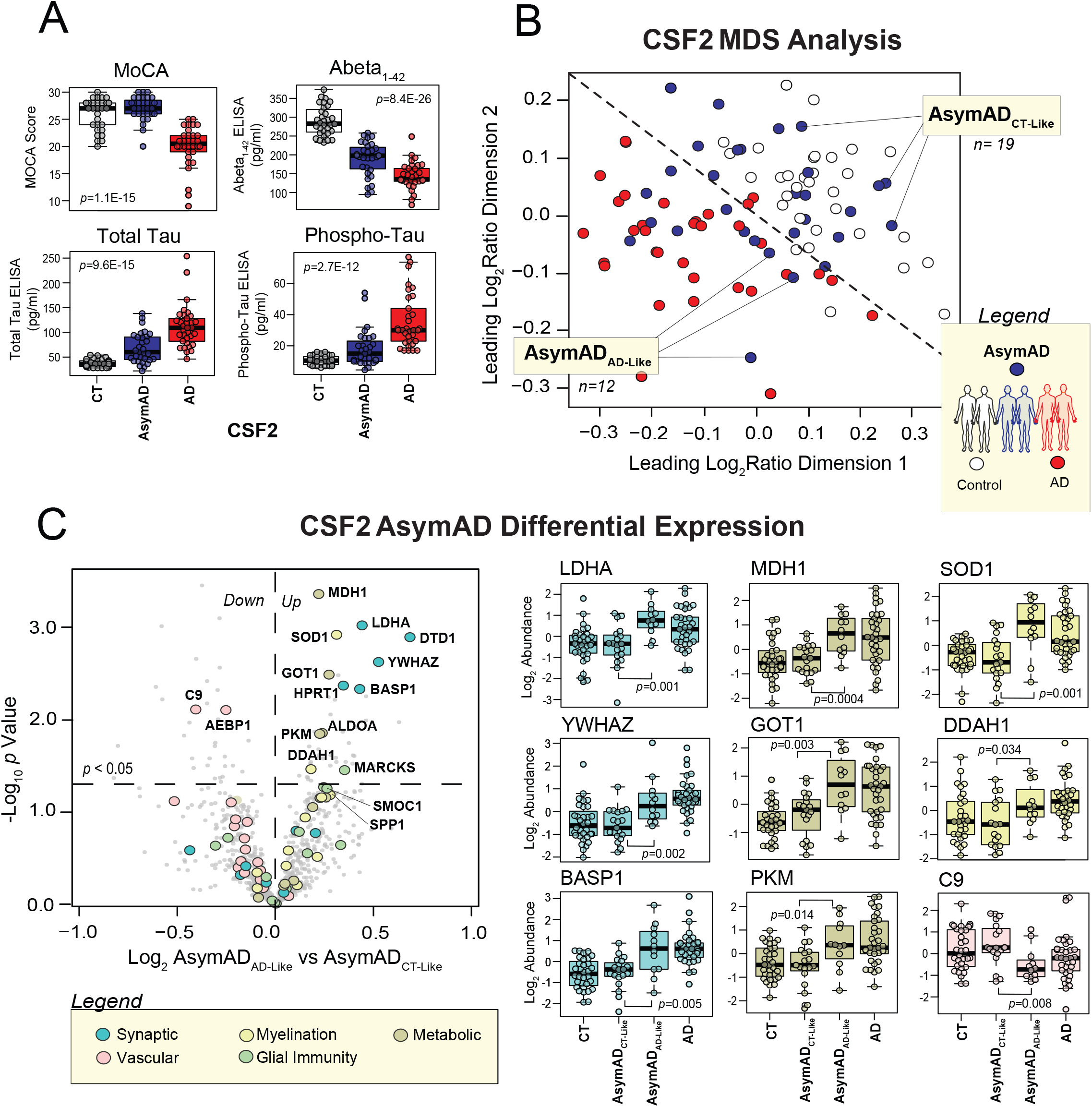
Validated CSF Biomarkers Classify Subgroups within AsymAD. A) MoCA scores and ELISA amyloid and tau levels plotted across the 96 cases of the CSF2 cohort. B) Multidimensional scaling analysis (MDS) of the 96 CSF2 cases based on the abundance levels of the 29 validated panel markers that demonstrated statistically significant AD alterations across both CSF datasets. This analysis yielded largely distinct clusters of control (white) and AD (red) cases. However, the AsymAD (blue) cases were scattered more variably throughout the plot. As shown, the MDS was used to subdivide the AsymAD group into two cohorts for subsequent analyses. One subgroup comprised those AsymAD cases that clustered more tightly with controls (*n*=19), while the other included those with marker profiles more closely resembling AD (*n*=12). C) Differential expression analysis between the proteomes of the 19 control-like and 12 AD-like AsymAD cases. On the left is a volcano plot displaying the log_2_ fold-change (x-axis) against the −log_10_ statistical *p* value for all CSF2 proteins differentially expressed between these two AsymAD groups. Enlarged colored data points represent the 70 validated panel markers. Each of the colors reflect a different panel; see figure legend. Overall, 14 of these validated markers (labeled) demonstrated significant differences between the two AsymAD subgroups. On the right, the abundance levels of 9 of these 14 markers are individually plotted across the 96 cases of the CSF2 cohort divided into control, control-like AsymAD, AD-like AsymAD, and AD groups. ANOVA with Tukey post-hoc correction was used to assess the statistical significance between the protein abundance changes observed between the two AsymAD subgroups. Abbreviations: CT, Control; AsymAD, Asymptomatic AD; AD, Alzheimer’s Disease; MoCA, Montreal Cognitive Assessment.

To determine the behavior of our validated panels in these AsymAD individuals, we applied a multidimensional scaling (MDS) analysis to all 96 of our CSF2 cases. MDS analysis allows for the visualization of similarities among cases based on certain variables in a dataset. To cluster cases, we used the abundance levels of the 29 most strongly validated markers from our five panels, i.e. those that demonstrated statistically significant AD abundance changes in both the CSF1 and CSF2 datasets (*p*<0.05). This MDS analysis demonstrated clear spatial clusters between control and AD cases (**Figure 7B**). However, the AsymAD cases were scattered with much more variability throughout the diagram. While certain AsymAD cases clustered unequivocally among the controls, others were situated on the nearly opposite end of the plot among the AD cases. Finally, a small group of AsymAD cases appeared to cluster near the center. In order to analyze this AsymAD heterogeneity in more detail, we used our MDS plot to define two groups among these preclinical cases. The first group comprised AsymAD cases that clustered closer to controls (*n*=19), while the second group harbored pre-symptomatic cases that more resembled the marker profiles of AD (*n*=12).

To identify those markers most capable of distinguishing these two AsymAD subgroups, we performed a differential expression analysis between the proteomes of the 19 control-like AsymAD and 12 AD-like AsymAD cases (**Table S9**). This volcano plot revealed 14 markers from our validated panels significantly altered between these two groups (**Figure 7C, left panel**). Most of these markers were members of the synaptic and metabolic panels (**Figure 7C, right panel**). However, SOD1 and the actin cross-linking protein MARCKS represented glial cell activity among this group of potential pre-clinical targets. The vascular panel also contributed two markers that were notably decreased in the AD-like AsymAD group, including the wound healing protein AEBP1 and complement family member C9. Interestingly, these 14 markers further highlighted the strong metabolic theme running throughout much of our validated panel, including elevated markers of neuronal metabolism (LDHA, HPRT1, MDH1, GOT1, PKM, ALDOA) and glial-mediated clearance of reactive oxygen species (SOD1, DDAH1). Overall, these results indicated that our validated panels may not only provide biomarkers relevant to symptomatic AD, but also yield markers potentially useful for the stratification and staging of asymptomatic disease.

## Discussion

The increasing recognition that a broad spectrum of pathologies contribute to AD and the repeated failures of amyloid-targeting drugs have highlighted the urgent need for biomarkers that more comprehensively reflect the complex mechanisms underlying this disease [4–6, 68]. While others have embarked on large-scale discovery-driven investigations of AD CSF, the context of emerging targets within the intricate landscape of brain-based pathophysiology has remained largely unexplored [61, 69]. To address this challenge, we applied an unbiased, integrative proteomics approach to the brain and CSF of AD subjects to identify fluid markers linked to a variety of processes in the diseased brain. Our results ultimately yielded five panels of brain-linked CSF markers that 1) reflected a wide variety of AD cortical pathophysiology, ranging from synaptic dysfunction to glial-mediated immunity, 2) demonstrated robust detectability across varied mass spectrometry platforms and reproducibility in independent analyses, and 3) harbored changes throughout both asymptomatic and symptomatic stages of disease. Overall, these findings represent a promising step toward the development of a systems-oriented biomarker tool capable of significantly advancing AD therapeutics. Our ability to map accessible fluid markers to brain-based cell types and functions could help meet many of the challenges facing AD drug development, from the identification of disease-modifying targets to the development of drug-monitoring markers reflective of meaningful target engagement. Furthermore, our validation studies revealed a potential role for our brain-linked CSF panels in the detailed risk stratification of individuals in the earliest presymptomatic stages of disease.

These findings strongly support the utility of data-driven network-based proteomics in the identification and clinical translation of AD biomarkers. Yet, in addition to its possible practical translations, this approach may also enhance our biological understanding of the critical protein systems governing disease pathogenesis. While our findings implicated a variety of protein modules in AD pathogenesis, a common theme of dysregulated energy metabolism emerged across all five of our validated marker panels. Metabolic proteins, such as HPRT1 and LDHA, were among those most strongly validated markers of our synaptic panel, demonstrating highly significant up-regulation in both CSF datasets. Furthermore, the validated vascular and glial panels also featured markers involved in the metabolism of oxidative species. These findings align with the critical role that metabolic processes play throughout the brain to meet the high energy demands of not only neurons, but also astrocytes and other glial cells [70]. In addition, our results support the growing evidence that altered redox potential and disrupted energy pathways may comprise the central link between several key processes implicated in AD pathogenesis, including mitochondrial dysregulation, glial-mediated inflammation, and vascular damage [49]. Furthermore, that such metabolic CSF biomarkers comprised the bulk of differentially abundant proteins between our control-like and AD-like AsymAD subjects suggests that these disruptions of redox potential and energy metabolism may be critical during the early preclinical stages of disease. Longitudinal studies will be necessary to further assess the performance of these biomarkers in disease progression.

The expression levels of our biomarker panels may also provide valuable biological insights. The divergent brain-CSF trends displayed by several of our marker panels are perhaps most interesting. One could refer to Aβ protein and its well-established divergence in AD brain and CSF. Its increased levels in brain as insoluble pools of peptide in plaques presumably results in decreased soluble peptide in the CSF. Yet, such an explanation fails to apply to the bulk of proteins found in our panels. Our neuron-associated synaptic and metabolic panels demonstrated the opposite expression pattern with decreased levels in the AD brain yet predominantly increased abundance in AD CSF. Given neurons are enriched with energy-generating mitochondria at synapses to fuel their numerous specialized signals [57], the parallels in expression of these two panels might be anticipated. However, it is unclear precisely why these panels demonstrated such divergent expression trends in the brain and CSF. Neuronal loss could account for the decreased abundance found in the demented brain but fails to explain the loss of synapse-associated proteins in the AsymAD brain, nor the increase in these synaptic proteins in the spinal fluid during both early and later stages of disease. One possible explanation is dysregulated endo- and exocytosis, both of which have been increasingly implicated in AD pathogenesis [71–75]. Indeed, aberrant synaptic glial-mediated phagocytosis of synapses, perhaps in response to aberrant intracellular Aβ production, has been proposed as a major contributor to the synaptic loss found early in disease [74]. The further processing of this synaptic pruning through exocytic pathways could then account for the increased release of synapse-associated proteins into the extracellular space. Indeed, multiple studies have demonstrated altered exosome content in the AD brain. Furthermore, aberrant exocytic pathways have even been linked to the secretion and propagation of Aβ and other pathogenic modules [76, 77]. Notably, the inhibition of exosome secretion may reduce AD-like pathology in a transgenic mouse model of AD [78]. Yet, while it is still widely debated whether exosome involvement is directly responsible for disease propagation, it may explain the increased levels of synaptic and metabolic proteins in AD CSF.

Meanwhile, the proteins comprising the vascular panel demonstrated a similar divergent expression pattern, featuring a modest increase in the AD brain but a stark decrease in levels in AD CSF. To explain these findings, one might look toward blood brain barrier (BBB) dysfunction. The BBB breakdown in AD has now been demonstrated by numerous independent postmortem human studies [48, 79]. These investigations have confirmed a variety of aberrant activity surrounding this tightly sealed layer of endothelial cells, including brain capillary leakages and the perivascular accumulation of blood-derived proteins [48]. This could provide a simple explanation for the increase of vascular proteins we identified in our Brain1 proteome. Yet, how does one explain the depletion of these same proteins in the spinal fluid? One possibility is that the brain is actively sequestering these molecules to address heightened levels of inflammation and oxidative stress. Several of the most decreased CSF1 proteins within this panel, particularly those involved in lipoprotein regulation, have been implicated in neuroprotective processes that suppress harmful levels of inflammation and reactive oxygen species. This is certainly true of paroxonase 1 (PON1), a lipoprotein-binding enzyme responsible for reducing oxidative stress levels in the circulation [65, 66]. AMBP, another significantly down-regulated vascular panel marker, serves as a precursor for the lipid transporter, bikunin, which has also been implicated in inflammation suppression and neuroprotection [80, 81].

The inability to directly probe biochemical disease mechanisms is often a limitation of discovery-driven proteomic analysis. Therefore, further studies are necessary to clarify the mechanisms underlying our panel levels. Future directions will also require investigation of these panels across other proteomic techniques, such as targeted methods like selective or parallel reaction monitoring. These studies should include analysis of not only our 70 validated markers, but also the nearly 300 brain-linked CSF biomarkers identified in our initial discovery-driven integrative approach. Such analyses could offer valuable additional insight into marker reproducibility and allow for panel optimization. The validation approach of the current study was designed to prioritize CSF targets that can be reliably measured by high-throughput proteomic techniques with minimal required sample preparation, hence increasing their potential for successful clinical translation. Indeed, using a FAIMS-based single shot proteomic approach in our validation step allowed us to forgo the time-consuming process of albumin depletion and dramatically reduced instrument time during the mass spectrometry analysis. Yet, this design ultimately sacrificed the depth of our validation proteome, preventing the validation of potentially relevant brain-linked fluid markers. Neurofilament light (NEFL) and neurogranin (NRGN) are two such examples of emerging AD biomarkers that mapped to overlapping modules during our integrative analysis but could not be detected using our high-throughput validation method. Therefore, employing other methods of validation (e.g. immunoassays) may prove informative.

Nevertheless, our integrative approach could still offer contextual insights regarding a range of promising biomarkers, regardless of their capacity for high-throughput validation. For instance, both NEFL and NRGN mapped to brain-derived networks highly consistent with their postulated roles in disease. NEFL’s link to the myelination panel supports its likely involvement in axonal degeneration, while NRGN’s ties to the synaptic panel, particularly the post-synaptic M12 module, are consistent with its known role in calmodulin-binding at the post-synaptic membrane and further implicates this protein in the synaptic degeneration critical to AD pathogenesis [20, 21, 82]. Overall, the current study offers a unique systems-based approach for the identification and validation of pathologically contextualized CSF AD biomarkers. The optimization of such marker panels across additional AD cohorts and proteomic platforms could prove promising for the advancement of AD risk stratification and therapeutics. In addition, future studies assessing the longitudinal behavior of these panels over time will be critical to determine which combination of markers best stratify risk in early disease and change in accordance to disease severity.

## Materials and Methods

### Sample Descriptions

#### Postmortem Brain Tissues

All brain tissues used in this study were derived from the dorsolateral prefrontal cortex and processed in the Emory Goizueta Alzheimer’s Disease Research Center (ADRC). Postmortem neuropathological evaluation of amyloid plaque distribution was performed according to the Consortium to Establish a Registry for Alzheimer’s Disease (CERAD) criteria [83], while the extent of neurofibrillary tangle pathology was assessed in accordance with the Braak staging system [84]. All AD cases met NIA-Reagan criteria for the diagnosis of AD (high likelihood) [85]. PD cases also met established criteria and guidelines for diagnosis [86]. Cases were classified as co-morbid AD and PD (AD/PD) when they met pathological criteria for amyloid plaque, neurofibrillary tangle, and Lewy body burden. Pathological and clinical evaluations for the asymptomatic AD (AsymAD) brain tissues were previously described [87]. Two cohorts of brain tissue were used in the proteomic studies. The Brain1 cohort included tissues from 10 healthy control, 10 PD, 10 AD/PD, and 10 AD cases. The Brain2 cohort included 19 cases identical to the Brain1 cohort (10 control and 9 AD cases), as well as 8 AsymAD cases unique to this cohort. All case metadata including disease state, gender, race, apolipoprotein (ApoE) genotype, age of death, Mini Mental State Examination (MMSE) and post-mortem interval (PMI) are provided in **Table 1**.

#### CSF Samples

All participants from whom CSF samples were collected provided informed consent under protocols approved by the Institutional Review Board (IRB) at Emory University. All patients received standardized cognitive assessments (including MoCA) in the Emory Cognitive Neurology clinic, the Emory Goizueta ADRC, and affiliated research studies (Emory Healthy Brain Study [EHBS] and Emory M^2^OVE-AD study). All diagnostic data were supplied by the ADRC and the Emory Cognitive Neurology Program. CSF was collected by lumbar puncture and banked according to 2014 ADC/NIA best practices guidelines (https://www.alz.washington.edu/BiospecimenTaskForce.html). For patients recruited from the Emory Cognitive Neurology Clinic, CSF samples were sent to Athena Diagnostics and assayed for Aβ_1-42_, Total Tau, and Phospho-Tau (CSF ADmark®) using the INNOTEST® assay platform. CSF samples collected from research participants in the ADRC, EHBS, and M^2^OVE-AD were assayed using the INNO-BIA AlzBio3 Luminex assay [88]. In total, there were two cohorts of CSF samples that were used in the proteomics studies. The CSF1 cohort contained samples from 20 healthy controls and 20 AD patients. CSF2 included spinal fluid obtained from three groups: healthy controls (*n*=32), AsymAD (*n*=31), and AD (*n*=33). AD and AsymAD cases were defined using established biomarker cutoff criteria for AD according to each assay platform [89, 90]. Cohort information is provided in **Table 1**.

### Multiplex Proteomics of CSF Samples

#### Immunodepletion and Digestion of CSF1 Samples

To increase the depth of our CSF1 dataset, immunodepletion of highly abundant proteins was employed prior to trypsin digestion. Briefly, 130 µL of CSF from each of the 40 individual CSF samples was incubated with equal volume (130µL) of High Select Top14 Abundant Protein Depletion Resin (Thermo Scientific, A36372) at room temperature in centrifuge columns (Thermo Scientific, A89868). After 15 minutes of rotation, the samples were centrifuged at 1,000g for 2 minutes. Sample flow-through was concentrated with a 3K Ultra Centrifugal Filter Device (Millipore, UFC500396) by centrifugation at 14,000g for 30 minutes. All sample volumes were diluted to 75 µL with phosphate buffered saline (PBS). Protein concentration and integrity was assessed by bicinchoninic acid (BCA) method according to manufacturer protocol (Thermo Scientific). Immunodepleted CSF (60 µL) from all 40 samples was digested with lysyl endopeptidase (LysC) and trypsin. Briefly, the samples were reduced and alkylated with 1.2 µL 0.5 M tris-2(-carboxyethyl)-phosphine (TCEP) and 3 µL 0.8M chloroacetamide (CAA) at 90°C for 10 minutes, followed by water bath sonication for 15 minutes. Samples were diluted with 193 µL 8M urea buffer (8 M urea, 100mM NaHPO4, pH 8.5) to a final concentration 6M urea. LysC (4.5µg, Wako) was used for overnight digestion at room temperature. Samples were then diluted to 1M urea with 50mM ammonium bicarbonate (ABC). An equal amount (4.5µg) of trypsin (Promega) was added and the samples subsequently incubated for 12 hours. The digested peptide solutions were acidified to a final concentration of 1% formic acid (FA) and 0.1% trifluoroacetic acid (TFA), followed by desalting with 100 mg C18 Sep-Pak columns (Waters) as described previously [16]. The peptides were subsequently eluted in 1 mL of 50% acetonitrile (ACN). In order to normalize protein quantification across batches [16], 100 μL aliquots from all 40 CSF samples were pooled to generate a pooled sample, which was then divided into 5 global internal standard (GIS) samples. All individual samples and the pooled standards were dried by speed vacuum (Labconco).

#### TMT Labeling of CSF1 Samples

All 40 samples and 5 GIS samples were divided into 5 batches, labeled using an 11-plex tandem mass tag (TMT) kit (Thermo Scientific, A34808, lot no. for TMT 10-plex: SI258088, 131C channel SJ258847), and derivatized as previously described [16]. See Data Availability section for sample to batch arrangement. Nine of the 11 TMT channels were utilized for labeling: 127N, 128N, 128C, 129N, 129C, 130N, 130C, 131N, 131C. Briefly, 5 mg of each TMT reagent was dissolved in 256 μL anhydrous ACN. Each CSF peptide digest was resuspended in 50 μL 100 mM triethylammonium bicarbonate (TEAB) buffer and 20.5 µl TMT reagent solution subsequently added. After 1 hour, the reaction was quenched with 4 μL 5% hydroxylamine (Thermo Scientific, 90115) for 15 minutes. After labeling, the peptide solutions were combined according to the batch arrangement. Each TMT batch was desalted with 100 mg C18 Sep-Pak columns (Waters) and dried by speed vacuum (Labconco).

#### High pH Fractionation of TMT-Labeled CSF1 Samples

High pH fractionation was performed as previously described [91] with slight modifications. The TMT-labeled peptides (160 ug) of CSF1 samples were dissolved in 100 μL of loading buffer (1mM ammonium formate, 2% (vol/vol) ACN), injected completely with an auto-sampler, and fractionated using a ZORBAX 300Extend-C18 column (2.1 mm x 150 mm, 3.5 µm, Agilent Technologies) on an Agilent 1100 HPLC system monitored at 280 nm. A total of 96 fractions were collected over a 60-min gradient of 100% mobile phase A (4.5mM ammonium formate (pH 10) in 2% vol/vol ACN) from 0-2 min, 0%–12% mobile phase B (4.5 mM ammonium formate (pH 10) in 90% vol/vol ACN) from 2-8 min, 12%–40% B from 8-36 min, 40%–44% B from 36-40 mins, 44%-60% B from 40-45 mins, and 60% B until completion with a flow rate of 0.4 mL/min. The 96 fractions were collected with an even time distribution and pooled into 30 fractions.

#### LC-MS/MS of TMT-Labeled CSF1 Samples

An equal volume of each of the 30 high-pH peptide fractions was resuspended in loading buffer (0.1% FA, 0.03% TFA, 1% ACN). Peptide eluents were separated on a self-packed C18 (1.9 µm Dr. Maisch, Germany) fused silica column (25 cm × 75μM internal diameter (ID), New Objective, Woburn, MA) by an Easy-nanoLC system (Thermo Scientific) and monitored on an Orbitrap HF-X mass spectrometer (Thermo Scientific). Elution was performed over a 120-min gradient at a rate of 225 nL/min with buffer B ranging from 1% to 90% (buffer A: 0.1% FA in water, buffer B: 0.1 % FA in ACN). The mass spectrometer was set to acquire data in positive ion mode using data-dependent acquisition. Each cycle consisted of one full MS scan followed by a maximum of 10 MS/MS scans. Full MS scans were collected at a resolution of 120,000 (400-1600 m/z range, 3×10^6^ AGC, 100 ms maximum ion injection time). All higher energy collision-induced dissociation (HCD) MS/MS spectra were acquired at a resolution of 45,000 (1.6 m/z isolation width, 35% collision energy, 1×10^5^ AGC target, 86 ms maximum ion time). Dynamic exclusion was set to exclude previously sequenced peaks for 20 seconds within a 10-ppm isolation window. See Data Availability section for all raw mass spectrometry files and search results.

#### Database Search and Protein Quantification of CSF1 Dataset

All raw files were analyzed using the Proteome Discoverer Suite (version 2.1, Thermo Scientific). MS/MS spectra were searched against the UniProtKB human proteome database (downloaded April 2015 with 90,411 total sequences). The Sequest HT search engine was used with the following parameters: fully tryptic specificity; maximum of two missed cleavages; minimum peptide length of 6; fixed modifications for TMT tags on lysine residues and peptide N-termini (+229.162932 Da) and carbamidomethylation of cysteine residues (+57.02146 Da); variable modifications for oxidation of methionine residues (+15.99492 Da) and deamidation of asparagine and glutamine (+0.984 Da); precursor mass tolerance of 20 ppm; and fragment mass tolerance of 0.05 Da. The Percolator node was used to filter peptide spectral matches (PSMs) to a false discovery rate (FDR) of less than 1%. Following spectral assignment, peptides were assembled into proteins and were further filtered based on the combined probabilities of their constituent peptides to a final FDR of 1%. In cases of redundancy, shared peptides were assigned to the protein sequence in adherence with the principles of parsimony. Reporter ions were quantified from MS2 scans using an integration tolerance of 20 ppm with the most confident centroid setting, as previously described [16].

#### Preparation of CSF2 Samples

In contrast to the CSF1 samples, those of CSF2 were not immunodepleted in preparation for mass spectrometry analysis. Instead, the protein concentrations for each of the 96 CSF2 samples were determined by BCA and then directly digested with LysC and trypsin. Briefly, 20 µL CSF from each sample was reduced and alkylated with 0.4 µL 0.5M TCEP and 2 µL 0.4M CAA at 90 °C for 10 minutes, followed by 15 minutes of water bath sonication. The samples were then further denatured in 67.2 µL of 8M urea (8M urea, 100mM NaHPO_4_, pH 8.5) to yield a final concentration of 6M urea and subsequently digested overnight with 1.9 µg LysC (1:10 enzyme to protein ratio). Following LysC digestion, the samples were diluted to 1M urea using 50mM ABC. An equivalent amount of trypsin (Promega) was then added (1:10 enzyme to protein ratio) and digestion was carried out for another 12 hours. After trypsin digestion, the peptide solutions were acidified with a 1% TFA and 10% FA solution to a final concentration of 0.1% TFA and 1% FA. Peptides were desalted with 100 mg C18 HLB columns (Waters) and eluted in 1 mL of 50% ACN. In order to normalize protein quantification across batches, 120 µL aliquots of eluted peptides from each of the 96 CSF samples were pooled to generate a GIS. All individual samples and the pooled standard were dried by speed vacuum (Labconco). To boost the signal of low abundance CSF proteins, a “boosting” sample (i.e. a biological sample mimicking study samples but accessible in a much larger quantity [92, 93]) was prepared by combining 125 µL from each of the 96 samples into a pooled CSF sample. This pooled sample was subsequently immunodepleted using 12 mL of High Select Top14 Abundant Protein Depletion Resin (Thermo Scientific, A36372), digested as described above, and included in the subsequent multiplex TMT assay.

#### TMT Labeling of CSF2 Samples

All 96 samples, as well as the GIS and boosting samples, were divided into 12 batches and labeled using an 11-plex TMT kit (Thermo Scientific, A34808, lot no. for TMT 10-plex: SI258088, 131C channel SJ258847). See Data Availability section for sample to batch arrangement. TMT labeling of these CSF2 samples was performed very similarly to the CSF1 samples with small alterations due to the inclusion of a boosting sample. Ten of the 11 TMT channels were utilized for labeling: 126 (boosting sample), 127N, 128N, 128C, 129N, 129C, 130N, 130C, 131N, 131C. Briefly, TMT reagent (5 mg) was dissolved in 256 μL anhydrous ACN. Each CSF peptide digest was resuspended in 50 μL 100 mM TEAB buffer and 20.5 µl TMT reagent solution subsequently added. The immunodepleted boosting sample was dissolved in 1.25 mL 100mM TEAB and labeled with 2×5mg of reagent (TMT 126 channel). For each plex, the pooled boosting channel was equivalent to 50-fold volume of each CSF sample. After 1 hour, these reactions were quenched with 8ul of 5% hydroxylamine for 15 minutes. After labeling, the peptide solutions were combined according to the batch arrangement. Each TMT batch was then desalted with 100 mg C18 Sep-Pak columns (Waters) and dried by speed vacuum (Labconco).

#### Single-shot FAIMS LC-MS/MS of TMT-Labeled CSF2 Samples

All samples were resuspended in equal volume of loading buffer (0.1% FA, 0.03% TFA, 1% ACN). Peptide eluents were separated on a self-packed C18 (1.9 µm, Dr. Maisch, Germany) fused silica column (25 cm × 75μM internal diameter (ID): New Objective, Woburn, MA) by an Easy-nLC system (Thermo Scientific) and monitored on an Orbitrap Fusion Lumos mass spectrometer (Thermo Scientific) interfaced with high-Field Asymmetric Waveform Ion Mobility Spectrometry (FAIMS). Sample elution was performed over a 180-min gradient with mobile phase A (0.1% of FA in water) and B (80% ACN in 0.1% FA) at a flow rate of 225 nL/min. The gradient was 1% to 8% B over 3 min, then 8% to 40% B over 160 min, then 40% to 99% B over 10 min, and finally 99% B for 10 min. The mass spectrometer was set to acquire data in positive ion mode using data-dependent acquisition and three (−50, −65 and −85) different compensation voltages (CV) [51]. Each of the three experiments consisted of a single CV and 1s cycles. Each cycle consisted of 1 full scan followed by as many MS2 and MS3 scans as possible within a 1s timeframe. Full MS scans were collected at a resolution of 120,000 (450-1500 m/z range, 4×10^5^ AGC, 50 ms maximum injection time). The collision induced dissociation (CID) MS/MS scans were collected in the ion trap with an isolation window of 0.7 m/z, collision energy of 35%, AGC setting of 1×10^4^, and a maximum injection time of 50 ms. The top 10 product ions were subjected to HCD synchronous precursor selection-based MS3 (SPS-MS3) as previously described [16]. For SPS-MS3 scans, the isolation window was set to 2 m/z, resolution to 50,000, AGC to 1×10^5^, and maximum injection time to 105 ms. A single preliminary run of TMT batch 1 using the above parameters was used to create a target inclusion list of peptides that specifically excluded those from the top 15 most abundant proteins. This inclusion list was used for all TMT batches. See Data Availability section of methods for all raw mass spectrometry files and search results.

#### Database Search and Protein Quantification of CSF2 Dataset

All raw files were analyzed using the Proteome Discoverer Suite (version 2.3, Thermo Scientific). MS/MS spectra were searched against the UniProtKB human proteome database (downloaded April 2015 with 90,411 total sequences). The Sequest HT search engine was used with the following parameters: fully tryptic specificity; maximum of two missed cleavages; minimum peptide length of 6; fixed modifications for TMT tags on lysine residues and peptide N-termini (+229.162932 Da) and carbamidomethylation of cysteine residues (+57.02146 Da); variable modifications for oxidation of methionine residues (+15.99492 Da), serine, threonine and tyrosine phosphorylation (+79.966 Da) and deamidation of asparagine and glutamine (+0.984 Da); precursor mass tolerance of 20 ppm; and a fragment mass tolerance of 0.6 Da. The Percolator node was used to filter PSMs to an FDR of less than 1%. Following spectral assignment, peptides were assembled into proteins and were further filtered based on the combined probabilities of their constituent peptides to a final FDR of 1%. In cases of redundancy, shared peptides were assigned to the protein sequence in adherence with the principles of parsimony. Reporter ions were quantified from MS3 scans using an integration tolerance of 20 ppm with the most confident centroid setting [16].

### Multiplex Proteomics of Brain Samples

#### Preparation of Brain Tissue Samples

The details regarding sample preparation of the Brain1 samples have been previously reported [16]. The Brain2 tissues were prepared and analyzed using very similar procedures. For each sample, approximately 100 mg of wet brain tissue was homogenized in 8M urea lysis buffer (8M urea, 10mM tris(hydroxymethyl) aminomethane, 100 mM NaHPO_4_, pH 8.5) containing Halt Protease and Phosphatase Inhibitor Cocktail (Thermo Scientific, 78440). Lysis buffer solution (500 μL) and stainless steel beads (∼100 μL, 0.9 to 2.0 mm blend, Next Advance) were added to individual RINO microcentrifuge tubes (Next Advance). Brain tissues were subsequently added immediately following excision. The samples were then placed into a Bullet Blender (Next Advance) in 4 °C cold room and homogenized for 2 full 5 min cycles. The lysates were transferred to new Eppendorf LoBind tubes and sonicated for 3 cycles, each consisting of 5 seconds of active sonication at 30% amplitude followed by 15 seconds on ice. The samples were subsequently centrifuged for 5 min at 15,000 g and the supernatant transferred to new tubes. BCA was used to determine the protein concentration for each sample prior to subsequent digestion. Approximately 100 μg of protein was digested for each Brain1 sample, while 500 μg of protein was digested for each Brain2 sample. Just prior to digestion, all samples were reduced with 1mM dithiothreitol (DTT) at room temperature for 30 minutes and alkylated with 5 mM iodoacetamide (IAA) in the dark for another 30 minutes. Lysyl endopeptidase (Wako) at 1:100 (w/w) was then added to each sample and digestion performed overnight. Samples were then diluted seven-fold with 50mM ABC. Trypsin (Promega) was then added at 1:50 (w/w) and digestion continued for another 16 hours. Each peptide solution was acidified to a final concentration of 1% (vol/vol) FA and 0.1% (vol/vol) TFA. The Brain1 samples were desalted with 100 mg C18 Sep-Pak columns (Waters) and eluted in 1 mL of 50% (vol/vol) ACN as described previously [16]. A 200 μL aliquot was removed from each Brain1 sample and combined to generate a pooled sample, which was subsequently divided into 10 GIS samples. Because a greater amount of protein was digested in the Brain2 analysis, each sample was desalted with a 200g Sep-Pak column and eluted in 3 mL of 50% (vol/vol) ACN. A 600 μL aliquot was removed from each Brain2 sample and combined to generate a pooled sample, which was then divided into 6 GIS samples. All digested peptide solutions from both Brain1 and Brain2 were dried by speed vacuum.

#### TMT Labeling of Brain Samples

All details of the Brain1 TMT labeling were previously reported [16]. In similar fashion, the 27 individual and 6 GIS samples of Brain2 were randomized into three batches and labeled using 11-plex TMT reagents. See Data Availability section for sample to batch arrangement. All 11 TMT channels were utilized for labeling. TMT reagent (5 mg) was dissolved in 56 μL anhydrous ACN. Due to the larger amount of Brain2 protein digested, two channels of 5 mg TMT reagent were combined to label each peptide solution. Each peptide solution was then reconstituted in 400 μL 100mM ‘TEAB buffer and 164 μL (3.2 mg) of labeling reagent subsequently added. After 1 hour, the reaction was quenched with 32 μL of 5% hydroxylamine. After labeling, the peptide solutions were combined according to the batch arrangement. Each TMT batch was then desalted with 500 mg C18 Sep-Pak columns (Waters) and eluted peptides were dried by speed vacuum (Labconco).

#### High pH Fractionation of TMT-Labeled Brain2 Samples

All TMT-labeled samples of Brain1 were subjected to electrostatic repulsion-hydrophilic interaction chromatography (ERLIC) fractionation prior to proteomic analysis as previously reported [16]. In contrast, the Brain2 samples were separated via high pH fractionation. For each Brain2 sample, approximately 4 mg of TMT-labeled peptides were resuspended in 850 μL loading buffer (1mM ammonium formate, 2% (vol/vol) ACN), injected completely with an auto-sampler, and fractionated using a ZORBAX 300Extend-C18 column (4.6 mm x 250 mm, 5 µm, Agilent Technologies) on an Agilent 1100 HPLC system monitored at 280 nm. A total of 96 fractions were collected over a 96-min gradient of 100% mobile phase A (4.5mM ammonium formate (pH 10) in 2% vol/vol acetonitrile) from 0-7 min, 0%–16% mobile phase B (4.5mM ammonium formate (pH 10) in 90% vol/vol acetonitrile) from 7-13 min, 16%–40% B from 13-73 min, 40%– 44% from 73-77 min, 44%-60% B from 77-82 mins, and 60% B until completion with a flow rate of 0.8 mL/min. The 96 fractions were collected with an even time distribution and pooled into 24 fractions.

#### LC-MS/MS of TMT-Labeled Brain2 Samples

Details of the mass spectrometry analysis of Brain1 samples were previously reported [16]. For the Brain2 cases, an equal volume of each of the 24 high-pH peptide fractions was resuspended in loading buffer (0.1% FA, 0.03% TFA, 1% ACN), and peptide eluents were separated on a self-packed C18 (1.9 um Dr. Maisch, Germany) fused silica column (25 cm × 75 μM internal diameter (ID), New Objective, Woburn, MA) by an Easy-nanoLC system (Thermo Scientific) and monitored on an Orbitrap Fusion mass spectrometer (Thermo Scientific). Elution was performed over a 140-min gradient at a rate of 225 nL/min with buffer B ranging from 1% to 90% (buffer A: 0.1% FA in water, buffer B: 80% ACN in water and 0.1% FA). The mass spectrometer was set to acquire data in top speed mode with 3-second cycles. Full MS scans were collected at a resolution of 120,000 (375-1500 m/z range, 4×10^5^ AGC, 50 ms maximum ion time). All HCD MS/MS spectra were acquired at a resolution of 50,000 (0.7 m/z isolation width, 38% collision energy, 1×10^5^ AGC target, 105 ms maximum ion time). Dynamic exclusion was set to exclude previously sequenced peaks for 20 sec within a 10-ppm isolation window. Only charge states from 2+ to 7+ were chosen for tandem MS/MS. See Data Availability section for all raw mass spectrometry files and search results.

#### Database Search and Protein Quantification of the Brain2 Dataset

The Brain1 dataset was searched and quantified as previously reported [16]. Raw data files from Orbitrap Fusion were processed using Proteome Discover Suite (version 2.1). MS/MS spectra were searched against the UniProtKB human proteome database (downloaded April 2015 with 90,411 total sequences). The Sequest HT search engine was used with the following parameters: fully tryptic specificity; maximum of two missed cleavages; minimum peptide length of 6; fixed modifications for TMT tags on lysine residues and peptide N-termini (+229.162932 Da) and carbamidomethylation of cysteine residues (+57.02146 Da); variable modifications for oxidation of methionine residues (+15.99492 Da) and deamidation of asparagine and glutamine (+0.984 Da); precursor mass tolerance of 20 ppm; and a fragment mass tolerance of 0.05 Da. The Percolator node was used to filter PSMs to an FDR of less than 1% using a target-decoy strategy. Following spectral assignment, peptides were assembled into proteins and were further filtered based on the combined probabilities of their constituent peptides to a final FDR of 1%. In cases of redundancy, shared peptides were assigned to the protein sequence in adherence with the principles of parsimony. Reporter ions were quantified from MS2 scans using an integration tolerance of 20 ppm with the most confident centroid setting [16].

### Data Analyses

#### Adjustment for Batch and Other Sources of Variance

For all four cohorts, only those proteins quantified in ≥50% of samples were included in the data analysis. Prior to subsequent analysis, all cohorts were also subjected to iterative outlier removal, which was performed as previously published [94] using Oldham’s ‘SampleNetworks’ v1.06 algorithm [17]. Sample connectivity (z score) greater than 3 standard deviations (SD) from the mean was used as the outlier criterion. This algorithm was applied repeatedly until no more outliers at this threshold could be found. In CSF1, one severe outlier was detected and therefore removed from all subsequent analyses. In contrast, no outliers were found among the CSF2 cohort. This algorithm also failed to detect outliers among the Brain1 and Brain2 datasets, even with the fold SD cutoff set more stringently to 2.5. For the CSF cohorts, batch correction was performed using a median polish algorithm for removing technical variance (i.e. variance due to tissue collection, cohort, or batch effects), derived from a two-way abundance-sample data table originally described by Tukey [95]. More detail on this method of batch correction can be found on GitHub (https://www.github.com/edammer/TAMPOR/). The brain datasets were not subjected to batch correction, though their protein abundance values were normalized by the GIS within each batch. Bootstrap regression for age at death, sex, and post-mortem interval (in the case of brain tissues) was performed on the protein log_2_ abundance ratios of all four cohorts. A principal component analysis (PCA) of the expression data confirmed appropriate regression of selected traits, using both the ‘SampleNetworks’ graphical output and R statistics for top five PC rank-based correlation to traits.

#### Differential Expression Analysis

Pairwise differentially expressed proteins were identified using student t-test followed by Benjamini-Hochberg (BH) FDR correction. Differential expression across three or more groups was performed using a one-way ANOVA followed by Tukey’s post-hoc test. Differential expression was presented in volcano plots, which were generated with the ggplot2 package in R v3.5.2.

#### Weighted Protein Correlation Network Analysis (WPCNA)

As previously described [10], a weighted Brain1 cohort protein co-expression network was derived from the post-regressed protein abundance values using the blockwiseModules WGCNA function (WGCNA 1.66 R package) with the following settings: soft threshold power beta=7.5, deepSplit=4, minimum module size=25, merge cut height=0.12, signed network with partitioning about medoids (PAM) respecting the dendrogram, TOMDenominator=”mean”, and a reassignment threshold of p<0.05. Hierarchical protein clustering analysis by protein expression pattern was performed within the WGCNA algorithm to generate the module dendrogram. Briefly, pairwise biweight midcorrelations were calculated between each protein pair during calculation of the signed adjacency matrix [96]. The connection strengths of multiple components within this matrix were then used to calculate a topological overlap matrix, representing measurements of protein expression pattern within the network [97]. Clustering was subsequently performed using the height similarity across all samples constructed via the pairwise correlations for all proteins within 1-TOM. Initial module identifications were established using dynamic tree cutting as implemented in the WGCNA::blockwiseModules function [98]. Module eigenproteins were defined as previously described [10], each representing the largest principle component of co-variance of all proteins within the module [99]. Pearson correlations between module eigenproteins and each quantified protein in Brain1 were performed and used to assign a measure of intra-module membership to each protein defined as kME. Module eigenproteins were also correlated with AD diagnosis and levels of amyloid and tau burden using biweight midcorrelation analysis.

#### Gene Ontology and Cell Type Enrichment Analyses

To characterize groups of proteins based on gene ontology annotation, we used GO Elite v1.2.5 as previously published [10, 94] with pruned output Fisher exact overrepresentation z scores visualized using R. The background proteome consisted of all proteins in each specific data set (*n*=2,875 for CSF1; *n*=792 for CSF2; *n*=8817 for Brain1; *n*=11244 for Brain2). Cell type enrichment for each of the modules in Brain1 was investigated as previously published [10, 94]. Briefly, the corresponding gene symbols of each module were cross-referenced with lists of genes known to be preferentially expressed in different cell types [100, 101]. Significance of cell type enrichment within each module was then determined using a one-tailed Fisher exact test (FET) and corrected for multiple comparisons by the BH FDR method.

#### Module Preservation Analysis

In order to assess Brain1 module preservation in Brain2, a co-expression network was also built for this cohort using similar WGCNA parameters with only slight modifications: soft threshold beta=15, deepSplit=3, and merge cut height=0.25. Preservation of the Brain1 modules in this Brain2 network was then tested using the R WGCNA::modulePreservation function with 500 permutations as previously described [10, 11].

#### Integrative Analysis of Brain and CSF Proteomic Datasets

Quantified CSF1 proteins were assessed for overrepresentation in Brain1 modules using a hypergeometric FET and those modules with BH-corrected p values of < 0.05 were considered significant. In similar fashion, separate FET analyses were performed to identify modules with significant overlap among differentially expressed CSF1 proteins, including those significantly up-regulated or down-regulated in AD compared to controls (p(CT-AD<0.05). The 15 modules with meaningful levels of CSF overlap from these three FET analyses were included in the five biomarker panels that were subsequently validated in the CSF2 cohort. CSF proteins were considered validated if they demonstrated altered levels in the same direction in both CSF1 and CSF2. In CSF2, composite z scores for the five panels of validated brain-linked CSF markers (n=70) were calculated and Kruskal-Wallis nonparametric ANOVA was performed across control, AsymAD, and AD groups.

#### Multidimensional Scaling Analysis

The multidimensional scaling (MDS) analysis of CSF2 data was performed using the limma package of R statistical software. A linear equation was selected to elicit a line across that the MDS plot separating control cases from those with AD. This line also divided the AsymAD cases into control-like and AD-like subgroups. All proteins measured in theses CSF2 AsymAD samples were then subjected to a t-test comparing their abundance levels in these subclasses of AsymAD.

## Supplementary Materials

Figure S1. Brain Proteome Highlights Disease-Specific Protein Alterations in AD

Figure S2. Analysis of AsymAD Brain Proteome Demonstrates Module Preservation and Presymptomatic Protein Alterations

Table S1. CSF1 Differential Expression by T-Test Analysis

Table S2. Top Gene Ontology (GO) Terms for Significantly Altered Proteins of CSF1

Table S3. Brain1 Differential Expression by ANOVA with Post-hoc Tukey Pairwise Comparisons

Table S4. Brain1 Module Assignments by Protein Correlations to Module Eigenproteins (kME)

Table S5. Top Gene Ontology (GO) Terms for Co-Expression Modules of Brain1

Table S6. Brain2 Differential Expression by ANOVA with Post-hoc Tukey Pairwise Comparisons

Table S7. Brain-Linked CSF Markers of Pre-Validation Panels

Table S8. CSF2 Differential Expression by ANOVA with Post-hoc Tukey Pairwise Comparisons

Table S9. CSF2 AsymAD Differential Expression by T-Test Analysis

## Funding

Support for this research was provided by funding from the National Institute on Aging (R01AG053960, R01AG057911, R01AG061800, RF1AG057471, RF1AG057470, R01AG061800, R01AG057911, R01AG057339), the Accelerating Medicine Partnership for AD (U01AG046161 and U01AG061357), the Emory Alzheimer’s Disease Research Center (P50AG025688), and the NINDS Emory Neuroscience Core (P30NS055077). Support was also provided by the Foundation for the National Institutes of Health (FNIH). N.T.S was also supported in part by grants from the Alzheimer’s Association (ALZ), Alzheimer’s Research UK (ARUK), The Michael J. Fox Foundation for Parkinson’s Research (MJFF), and the Weston Brain Institute (11060).

## Author Contributions

Conceptualization, L.H., E.B.D., E.C.B.J., J.J.L., A.I.L., and N.T.S.; Methodology, L.H., L.P., E.B.D., D.M.D., and N.T.S.; Investigation, L.H., L.P., E.B.D., D.M.D., M.Z., and N.T.S.; Formal Analysis, E.B.D.; Writing – Original Draft, L.H., L.P., E.B.D.; Writing – Review & Editing, L.H., E.C.B.J., J.J.L., A.I.L, and N.T.S.; Funding Acquisition, A.I.L and N.T.S.; Resources, M.G., I.H., J.J.L.; Supervision, A.I.L., J.J.L., and N.T.S.

## Competing Interests

The authors declare no conflicts of interest.

## Data Availability

All raw proteomic data generated contributing to the described work will be deposited electronically on SYNAPSE (https://www.synapse.org/#!Synapse:syn20821165/wiki/596086). The results published here are in whole or in part based on data obtained from the AMP-AD Knowledge Portal (https://adknowledgeportal.synapse.org).

---

**Figure S1.**
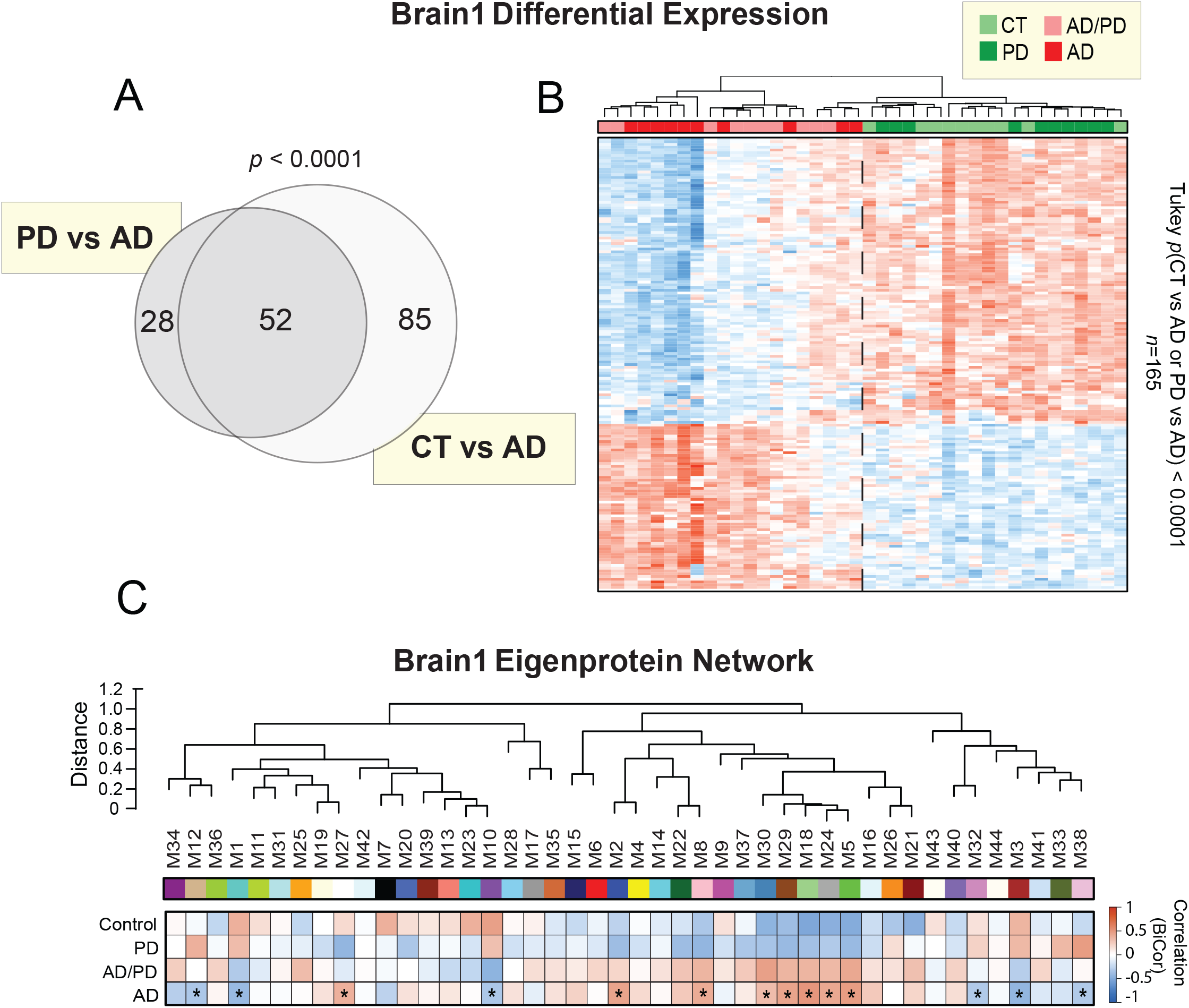
Brain1 Proteome Highlights Disease-Specific Protein Alterations in AD. A) Venn diagram of the 165 most highly altered proteins (*p*<0.0001) over two pairwise comparisons of the Brain1 proteome (PD vs AD, CT vs AD). There were no proteins with such highly significant changes across a comparison of controls and PD. B) Supervised cluster analysis across the 40 cases of the Brain1 cohort using these 165 highly altered proteins. This analysis resulted in the successful distinction between cases harboring AD pathology (AD=10, AD/PD=10) and amyloid-free control and PD cases. C) Biweight midcorrelation (bicor) analysis of the 44 co-expression modules of the Brain1 proteome to disease status. Asterisks highlight modules with statistically significant (*p*<0.05) correlations to AD status. Abbreviations: CT, Control; PD, Parkinson’s disease; AD, Alzheimer’s disease.

**Figure S2.**
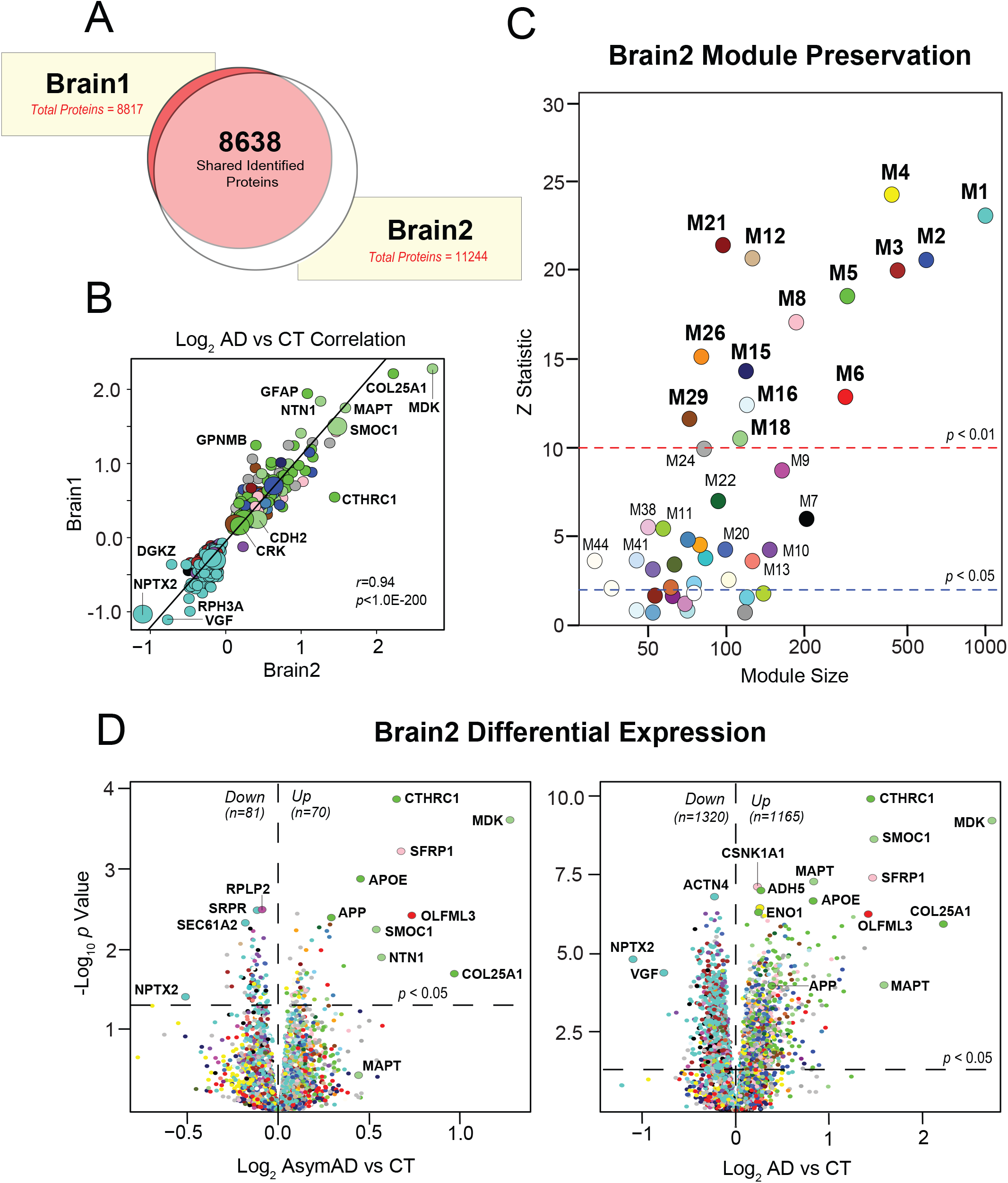
Brain2 Proteome Demonstrates Module Preservation Across Asymptomatic AD Cases. A) Venn diagram highlighting the high degree of overlap between proteins identified in the Brain1 and Brain2 proteomes. B) Correlation analysis of the differential abundance (log_2_ AD vs CT) in Brain1 and Brain2 of proteins that demonstrated significantly altered levels in both proteomes (*n*=450). Data points are colored by Brain1 module membership. Among proteins up-regulated in AD, those most highly correlated between the two datasets were members of the glial-enriched M5 (green) and M18 (light green) modules. Among proteins down-regulated in AD, those with the most concordant changes largely represented the synapse-associated M1 (turquoise) module. Pearson correlation coefficient and *p* value were used to assess the degree of correlation. C) Preservation analysis of the Brain1 modules in the Brain2 dataset. Of the 44 Brain1 co-expression modules, 36 were significantly preserved in the Brain2 proteome, with z-summary scores > 2 (*p*<0.05, above blue line). Fourteen modules were exceptionally preserved (*p*<0.01, above red line) in the Brain2 dataset. D) Volcano plots displaying the log_2_ fold-change (x-axis) against the −log_10_ statistical *p* value for all proteins differentially expressed in the Brain2 proteome between pairwise comparisons of control vs AsymAD cases (left) and control vs AD cases (right). Data points are colored by Brain1 module membership. Abbreviations: CT, Control; AsymAD, Asymptomatic AD; AD, Alzheimer’s Disease; M, Module.

## Notes

https://www.synapse.org/#!Synapse:syn20821165/wiki/596086

## References

1. Henry, M.S., A.P. Passmore, S. Todd, B. McGuinness, D. Craig, and J.A. Johnston, The development of effective biomarkers for Alzheimer’s disease: a review. Int J Geriatr Psychiatry, 2013. 28(4): p. 331–40.

2. Blennow, K., H. Hampel, M. Weiner, and H. Zetterberg, Cerebrospinal fluid and plasma biomarkers in Alzheimer disease. Nat Rev Neurol, 2010. 6(3): p. 131–44.

3. Jack, C.R., Jr., D.A. Bennett, K. Blennow, M.C. Carrillo, B. Dunn, S.B. Haeberlein, D.M. Holtzman, W. Jagust, F. Jessen, J. Karlawish, E. Liu, J.L. Molinuevo, T. Montine, C. Phelps, K.P. Rankin, C.C. Rowe, P. Scheltens, E. Siemers, H.M. Snyder, R. Sperling, and Contributors, NIA-AA Research Framework: Toward a biological definition of Alzheimer’s disease. Alzheimers Dement, 2018. 14(4): p. 535–562.

4. Lista, S., H. Zetterberg, S.E. O’Bryant, K. Blennow, and H. Hampel, Evolving Relevance of Neuroproteomics in Alzheimer’s Disease. Methods Mol Biol, 2017. 1598: p. 101–115.

5. Castrillo, J.I., S. Lista, H. Hampel, and C.W. Ritchie, Systems Biology Methods for Alzheimer’s Disease Research Toward Molecular Signatures, Subtypes, and Stages and Precision Medicine: Application in Cohort Studies and Trials. Methods Mol Biol, 2018. 1750: p. 31–66.

6. Boyle, P.A., L. Yu, S.E. Leurgans, R.S. Wilson, R. Brookmeyer, J.A. Schneider, and D.A. Bennett, Attributable risk of Alzheimer’s dementia attributed to age-related neuropathologies. Ann Neurol, 2019. 85(1): p. 114–124.

7. McKhann, G.M., D.S. Knopman, H. Chertkow, B.T. Hyman, C.R. Jack, Jr., C.H. Kawas, W.E. Klunk, W.J. Koroshetz, J.J. Manly, R. Mayeux, R.C. Mohs, J.C. Morris, M.N. Rossor, P. Scheltens, M.C. Carrillo, B. Thies, S. Weintraub, and C.H. Phelps, The diagnosis of dementia due to Alzheimer’s disease: recommendations from the National Institute on Aging-Alzheimer’s Association workgroups on diagnostic guidelines for Alzheimer’s disease. Alzheimers Dement, 2011. 7(3): p. 263–9.

8. Dai, J., E.C.B. Johnson, E.B. Dammer, D.M. Duong, M. Gearing, J.J. Lah, A.I. Levey, T.S. Wingo, and N.T. Seyfried, Effects of APOE Genotype on Brain Proteomic Network and Cell Type Changes in Alzheimer’s Disease. Front Mol Neurosci, 2018. 11: p. 454.

9. Johnson, E.C.B., E.B. Dammer, D.M. Duong, L. Yin, M. Thambisetty, J.C. Troncoso, J.J. Lah, A.I. Levey, and N.T. Seyfried, Deep proteomic network analysis of Alzheimer’s disease brain reveals alterations in RNA binding proteins and RNA splicing associated with disease. Mol Neurodegener, 2018. 13(1): p. 52.

10. Seyfried, N.T., E.B. Dammer, V. Swarup, D. Nandakumar, D.M. Duong, L. Yin, Q. Deng, T. Nguyen, C.M. Hales, T. Wingo, J. Glass, M. Gearing, M. Thambisetty, J.C. Troncoso, D.H. Geschwind, J.J. Lah, and A.I. Levey, A Multi-network Approach Identifies Protein-Specific Co-expression in Asymptomatic and Symptomatic Alzheimer’s Disease. Cell Syst, 2017. 4(1): p. 60–72 e4.

11. Umoh, M.E., E.B. Dammer, J. Dai, D.M. Duong, J.J. Lah, A.I. Levey, M. Gearing, J.D. Glass, and N.T. Seyfried, A proteomic network approach across the ALS-FTD disease spectrum resolves clinical phenotypes and genetic vulnerability in human brain. EMBO Mol Med, 2018. 10(1): p. 48–62.

12. Wingo, A.P., E.B. Dammer, M.S. Breen, B.A. Logsdon, D.M. Duong, J.C. Troncosco, M. Thambisetty, T.G. Beach, G.E. Serrano, E.M. Reiman, R.J. Caselli, J.J. Lah, N.T. Seyfried, A.I. Levey, and T.S. Wingo, Large-scale proteomic analysis of human brain identifies proteins associated with cognitive trajectory in advanced age. Nat Commun, 2019. 10(1): p. 1619.

13. Higginbotham, L., E.B. Dammer, D.M. Duong, E. Modeste, T.J. Montine, J.J. Lah, A.I. Levey, and N.T. Seyfried, Network Analysis of a Membrane-Enriched Brain Proteome across Stages of Alzheimer’s Disease. Proteomes, 2019. 7(3).

14. Roche, S., A. Gabelle, and S. Lehmann, Clinical proteomics of the cerebrospinal fluid: Towards the discovery of new biomarkers. Proteomics Clin Appl, 2008. 2(3): p. 428–36.

15. Rauniyar, N. and J.R. Yates, 3rd, Isobaric labeling-based relative quantification in shotgun proteomics. J Proteome Res, 2014. 13(12): p. 5293–309.

16. Ping, L., D.M. Duong, L. Yin, M. Gearing, J.J. Lah, A.I. Levey, and N.T. Seyfried, Global quantitative analysis of the human brain proteome in Alzheimer’s and Parkinson’s Disease. Sci Data, 2018. 5: p. 180036.

17. Oldham, M.C., G. Konopka, K. Iwamoto, P. Langfelder, T. Kato, S. Horvath, and D.H. Geschwind, Functional organization of the transcriptome in human brain. Nat Neurosci, 2008. 11(11): p. 1271–82.

18. Olsson, B., R. Lautner, U. Andreasson, A. Ohrfelt, E. Portelius, M. Bjerke, M. Holtta, C. Rosen, C. Olsson, G. Strobel, E. Wu, K. Dakin, M. Petzold, K. Blennow, and H. Zetterberg, CSF and blood biomarkers for the diagnosis of Alzheimer’s disease: a systematic review and meta-analysis. Lancet Neurol, 2016. 15(7): p. 673–684.

19. Muszynski, P., M. Groblewska, A. Kulczynska-Przybik, A. Kulakowska, and B. Mroczko, YKL-40 as a Potential Biomarker and a Possible Target in Therapeutic Strategies of Alzheimer’s Disease. Curr Neuropharmacol, 2017. 15(6): p. 906–917.

20. Thorsell, A., M. Bjerke, J. Gobom, E. Brunhage, E. Vanmechelen, N. Andreasen, O. Hansson, L. Minthon, H. Zetterberg, and K. Blennow, Neurogranin in cerebrospinal fluid as a marker of synaptic degeneration in Alzheimer’s disease. Brain Res, 2010. 1362: p. 13–22.

21. Kester, M.I., C.E. Teunissen, D.L. Crimmins, E.M. Herries, J.H. Ladenson, P. Scheltens, W.M. van der Flier, J.C. Morris, D.M. Holtzman, and A.M. Fagan, Neurogranin as a Cerebrospinal Fluid Biomarker for Synaptic Loss in Symptomatic Alzheimer Disease. JAMA Neurol, 2015. 72(11): p. 1275–80.

22. Blennow, K. and H. Zetterberg, Biomarkers for Alzheimer’s disease: current status and prospects for the future. J Intern Med, 2018. 284(6): p. 643–663.

23. Bereczki, E., R.M. Branca, P.T. Francis, J.B. Pereira, J.H. Baek, T. Hortobagyi, B. Winblad, C. Ballard, J. Lehtio, and D. Aarsland, Synaptic markers of cognitive decline in neurodegenerative diseases: a proteomic approach. Brain, 2018. 141(2): p. 582–595.

24. Pottiez, G., L. Yang, T. Stewart, N. Song, P. Aro, D.R. Galasko, J.F. Quinn, E.R. Peskind, M. Shi, and J. Zhang, Mass-Spectrometry-Based Method To Quantify in Parallel Tau and Amyloid beta 1-42 in CSF for the Diagnosis of Alzheimer’s Disease. J Proteome Res, 2017. 16(3): p. 1228–1238.

25. Mawuenyega, K.G., T. Kasten, W. Sigurdson, and R.J. Bateman, Amyloid-beta isoform metabolism quantitation by stable isotope-labeled kinetics. Anal Biochem, 2013. 440(1): p. 56–62.

26. Langfelder, P. and S. Horvath, WGCNA: an R package for weighted correlation network analysis. BMC Bioinformatics, 2008. 9: p. 559.

27. Miller, J.A., M.C. Oldham, and D.H. Geschwind, A systems level analysis of transcriptional changes in Alzheimer’s disease and normal aging. J Neurosci, 2008. 28(6): p. 1410–20.

28. Miller, J.A., S. Horvath, and D.H. Geschwind, Divergence of human and mouse brain transcriptome highlights Alzheimer disease pathways. Proc Natl Acad Sci U S A, 2010. 107(28): p. 12698–703.

29. Voineagu, I., X. Wang, P. Johnston, J.K. Lowe, Y. Tian, S. Horvath, J. Mill, R.M. Cantor, B.J. Blencowe, and D.H. Geschwind, Transcriptomic analysis of autistic brain reveals convergent molecular pathology. Nature, 2011. 474(7351): p. 380–4.

30. Zhang, B., C. Gaiteri, L.G. Bodea, Z. Wang, J. McElwee, A.A. Podtelezhnikov, C. Zhang, T. Xie, L. Tran, R. Dobrin, E. Fluder, B. Clurman, S. Melquist, M. Narayanan, C. Suver, H. Shah, M. Mahajan, T. Gillis, J. Mysore, M.E. MacDonald, J.R. Lamb, D.A. Bennett, C. Molony, D.J. Stone, V. Gudnason, A.J. Myers, E.E. Schadt, H. Neumann, J. Zhu, and V. Emilsson, Integrated systems approach identifies genetic nodes and networks in late-onset Alzheimer’s disease. Cell, 2013. 153(3): p. 707–20.

31. Takeda, K. and S. Akira, Toll-like receptors in innate immunity. Int Immunol, 2005. 17(1): p. 1–14.

32. Liu, J., C. Qian, and X. Cao, Post-Translational Modification Control of Innate Immunity. Immunity, 2016. 45(1): p. 15–30.

33. Heneka, M.T., M.J. Carson, J. El Khoury, G.E. Landreth, F. Brosseron, D.L. Feinstein, A.H. Jacobs, T. Wyss-Coray, J. Vitorica, R.M. Ransohoff, K. Herrup, S.A. Frautschy, B. Finsen, G.C. Brown, A. Verkhratsky, K. Yamanaka, J. Koistinaho, E. Latz, A. Halle, G.C. Petzold, T. Town, D. Morgan, M.L. Shinohara, V.H. Perry, C. Holmes, N.G. Bazan, D.J. Brooks, S. Hunot, B. Joseph, N. Deigendesch, O. Garaschuk, E. Boddeke, C.A. Dinarello, J.C. Breitner, G.M. Cole, D.T. Golenbock, and M.P. Kummer, Neuroinflammation in Alzheimer’s disease. Lancet Neurol, 2015. 14(4): p. 388–405.

34. Desler, C., M.S. Lillenes, T. Tonjum, and L.J. Rasmussen, The Role of Mitochondrial Dysfunction in the Progression of Alzheimer’s Disease. Curr Med Chem, 2018. 25(40): p. 5578–5587.

35. Duits, F.H., G. Brinkmalm, C.E. Teunissen, A. Brinkmalm, P. Scheltens, W.M. Van der Flier, H. Zetterberg, and K. Blennow, Synaptic proteins in CSF as potential novel biomarkers for prognosis in prodromal Alzheimer’s disease. Alzheimers Res Ther, 2018. 10(1): p. 5.

36. Molinuevo, J.L., S. Ayton, R. Batrla, M.M. Bednar, T. Bittner, J. Cummings, A.M. Fagan, H. Hampel, M.M. Mielke, A. Mikulskis, S. O’Bryant, P. Scheltens, J. Sevigny, L.M. Shaw, H.D. Soares, G. Tong, J.Q. Trojanowski, H. Zetterberg, and K. Blennow, Current state of Alzheimer’s fluid biomarkers. Acta Neuropathol, 2018. 136(6): p. 821–853.

37. Querfurth, H.W. and F.M. LaFerla, Alzheimer’s disease. N Engl J Med, 2010. 362(4): p. 329–44.

38. Sperling, R.A., P.S. Aisen, L.A. Beckett, D.A. Bennett, S. Craft, A.M. Fagan, T. Iwatsubo, C.R. Jack, Jr., J. Kaye, T.J. Montine, D.C. Park, E.M. Reiman, C.C. Rowe, E. Siemers, Y. Stern, K. Yaffe, M.C. Carrillo, B. Thies, M. Morrison-Bogorad, M.V. Wagster, and C.H. Phelps, Toward defining the preclinical stages of Alzheimer’s disease: recommendations from the National Institute on Aging-Alzheimer’s Association workgroups on diagnostic guidelines for Alzheimer’s disease. Alzheimers Dement, 2011. 7(3): p. 280–92.

39. Bennett, D.A., J.A. Schneider, Z. Arvanitakis, J.F. Kelly, N.T. Aggarwal, R.C. Shah, and R.S. Wilson, Neuropathology of older persons without cognitive impairment from two community-based studies. Neurology, 2006. 66(12): p. 1837–44.

40. Troncoso, J.C., A.M. Cataldo, R.A. Nixon, J.L. Barnett, M.K. Lee, F. Checler, D.R. Fowler, J.E. Smialek, B. Crain, L.J. Martin, and C.H. Kawas, Neuropathology of preclinical and clinical late-onset Alzheimer’s disease. Ann Neurol, 1998. 43(5): p. 673–6.

41. Driscoll, I. and J. Troncoso, Asymptomatic Alzheimer’s disease: a prodrome or a state of resilience? Curr Alzheimer Res, 2011. 8(4): p. 330–5.

42. Nelson, P.T., I. Alafuzoff, E.H. Bigio, C. Bouras, H. Braak, N.J. Cairns, R.J. Castellani, B.J. Crain, P. Davies, K. Del Tredici, C. Duyckaerts, M.P. Frosch, V. Haroutunian, P.R. Hof, C.M. Hulette, B.T. Hyman, T. Iwatsubo, K.A. Jellinger, G.A. Jicha, E. Kovari, W.A. Kukull, J.B. Leverenz, S. Love, I.R. Mackenzie, D.M. Mann, E. Masliah, A.C. McKee, T.J. Montine, J.C. Morris, J.A. Schneider, J.A. Sonnen, D.R. Thal, J.Q. Trojanowski, J.C. Troncoso, T. Wisniewski, R.L. Woltjer, and T.G. Beach, Correlation of Alzheimer disease neuropathologic changes with cognitive status: a review of the literature. J Neuropathol Exp Neurol, 2012. 71(5): p. 362–81.

43. Brellier, F., S. Ruggiero, D. Zwolanek, E. Martina, D. Hess, M. Brown-Luedi, U. Hartmann, M. Koch, A. Merlo, M. Lino, and R. Chiquet-Ehrismann, SMOC1 is a tenascin-C interacting protein over-expressed in brain tumors. Matrix Biol, 2011. 30(3): p. 225–33.

44. Okada, I., H. Hamanoue, K. Terada, T. Tohma, A. Megarbane, E. Chouery, J. Abou-Ghoch, N. Jalkh, O. Cogulu, F. Ozkinay, K. Horie, J. Takeda, T. Furuichi, S. Ikegawa, K. Nishiyama, S. Miyatake, A. Nishimura, T. Mizuguchi, N. Niikawa, F. Hirahara, T. Kaname, K. Yoshiura, Y. Tsurusaki, H. Doi, N. Miyake, T. Furukawa, N. Matsumoto, and H. Saitsu, SMOC1 is essential for ocular and limb development in humans and mice. Am J Hum Genet, 2011. 88(1): p. 30–41.

45. Xiao, M.F., D. Xu, M.T. Craig, K.A. Pelkey, C.C. Chien, Y. Shi, J. Zhang, S. Resnick, O. Pletnikova, D. Salmon, J. Brewer, S. Edland, J. Wegiel, B. Tycko, A. Savonenko, R.H. Reeves, J.C. Troncoso, C.J. McBain, D. Galasko, and P.F. Worley, NPTX2 and cognitive dysfunction in Alzheimer’s Disease. Elife, 2017. 6.

46. Sathe, G., C.H. Na, S. Renuse, A.K. Madugundu, M. Albert, A. Moghekar, and A. Pandey, Quantitative Proteomic Profiling of Cerebrospinal Fluid to Identify Candidate Biomarkers for Alzheimer’s Disease. Proteomics Clin Appl, 2019. 13(4): p. e1800105.

47. Rader, D.J. and K.A. Dugi, The endothelium and lipoproteins: insights from recent cell biology and animal studies. Semin Thromb Hemost, 2000. 26(5): p. 521–8.

48. Nelson, A.R., M.D. Sweeney, A.P. Sagare, and B.V. Zlokovic, Neurovascular dysfunction and neurodegeneration in dementia and Alzheimer’s disease. Biochim Biophys Acta, 2016. 1862(5): p. 887–900.

49. Yin, F., H. Sancheti, I. Patil, and E. Cadenas, Energy metabolism and inflammation in brain aging and Alzheimer’s disease. Free Radic Biol Med, 2016. 100: p. 108–122.

50. Consensus report of the Working Group on: “Molecular and Biochemical Markers of Alzheimer’s Disease”. The Ronald and Nancy Reagan Research Institute of the Alzheimer’s Association and the National Institute on Aging Working Group. Neurobiol Aging, 1998. 19(2): p. 109–16.

51. Hebert, A.S., S. Prasad, M.W. Belford, D.J. Bailey, G.C. McAlister, S.E. Abbatiello, R. Huguet, E.R. Wouters, J.J. Dunyach, D.R. Brademan, M.S. Westphall, and J.J. Coon, Comprehensive Single-Shot Proteomics with FAIMS on a Hybrid Orbitrap Mass Spectrometer. Anal Chem, 2018. 90(15): p. 9529–9537.

52. Sun, Y., X.S. Yin, H. Guo, R.K. Han, R.D. He, and L.J. Chi, Elevated osteopontin levels in mild cognitive impairment and Alzheimer’s disease. Mediators Inflamm, 2013. 2013: p. 615745.

53. Rentsendorj, A., J. Sheyn, D.T. Fuchs, D. Daley, B.C. Salumbides, H.E. Schubloom, N.J. Hart, S. Li, E.Y. Hayden, D.B. Teplow, K.L. Black, Y. Koronyo, and M. Koronyo-Hamaoui, A novel role for osteopontin in macrophage-mediated amyloid-beta clearance in Alzheimer’s models. Brain Behav Immun, 2018. 67: p. 163–180.

54. Comi, C., M. Carecchio, A. Chiocchetti, S. Nicola, D. Galimberti, C. Fenoglio, G. Cappellano, F. Monaco, E. Scarpini, and U. Dianzani, Osteopontin is increased in the cerebrospinal fluid of patients with Alzheimer’s disease and its levels correlate with cognitive decline. J Alzheimers Dis, 2010. 19(4): p. 1143–8.

55. Goetzl, E.J., D. Kapogiannis, J.B. Schwartz, I.V. Lobach, L. Goetzl, E.L. Abner, G.A. Jicha, A.M. Karydas, A. Boxer, and B.L. Miller, Decreased synaptic proteins in neuronal exosomes of frontotemporal dementia and Alzheimer’s disease. FASEB J, 2016. 30(12): p. 4141–4148.

56. De Vos, A., D. Jacobs, H. Struyfs, E. Fransen, K. Andersson, E. Portelius, U. Andreasson, D. De Surgeloose, D. Hernalsteen, K. Sleegers, C. Robberecht, C. Van Broeckhoven, H. Zetterberg, K. Blennow, S. Engelborghs, and E. Vanmechelen, C-terminal neurogranin is increased in cerebrospinal fluid but unchanged in plasma in Alzheimer’s disease. Alzheimers Dement, 2015. 11(12): p. 1461–1469.

57. Ly, C.V. and P. Verstreken, Mitochondria at the synapse. Neuroscientist, 2006. 12(4): p. 291–9.

58. Keating, D.J., Mitochondrial dysfunction, oxidative stress, regulation of exocytosis and their relevance to neurodegenerative diseases. J Neurochem, 2008. 104(2): p. 298–305.

59. Zhao, C., T. Li, B. Han, W. Yue, L. Shi, H. Wang, Y. Guo, and Z. Lu, DDAH1 deficiency promotes intracellular oxidative stress and cell apoptosis via a miR-21-dependent pathway in mouse embryonic fibroblasts. Free Radic Biol Med, 2016. 92: p. 50–60.

60. Bunton-Stasyshyn, R.K., R.A. Saccon, P. Fratta, and E.M. Fisher, SOD1 Function and Its Implications for Amyotrophic Lateral Sclerosis Pathology: New and Renascent Themes. Neuroscientist, 2015. 21(5): p. 519–29.

61. Saccon, R.A., R.K. Bunton-Stasyshyn, E.M. Fisher, and P. Fratta, Is SOD1 loss of function involved in amyotrophic lateral sclerosis? Brain, 2013. 136(Pt 8): p. 2342–58.

62. Lu, S.C., Glutathione synthesis. Biochim Biophys Acta, 2013. 1830(5): p. 3143–53.

63. Bandopadhyay, R., A.E. Kingsbury, M.R. Cookson, A.R. Reid, I.M. Evans, A.D. Hope, A.M. Pittman, T. Lashley, R. Canet-Aviles, D.W. Miller, C. McLendon, C. Strand, A.J. Leonard, P.M. Abou-Sleiman, D.G. Healy, H. Ariga, N.W. Wood, R. de Silva, T. Revesz, J.A. Hardy, and A.J. Lees, The expression of DJ-1 (PARK7) in normal human CNS and idiopathic Parkinson’s disease. Brain, 2004. 127(Pt 2): p. 420–30.

64. Choi, D.J., J.H. Eun, B.G. Kim, I. Jou, S.M. Park, and E.H. Joe, A Parkinson’s disease gene, DJ-1, repairs brain injury through Sox9 stabilization and astrogliosis. Glia, 2018. 66(2): p. 445–458.

65. Aluganti Narasimhulu, C., C. Mitra, D. Bhardwaj, K.Y. Burge, and S. Parthasarathy, Alzheimer’s Disease Markers in Aged ApoE-PON1 Deficient Mice. J Alzheimers Dis, 2019. 67(4): p. 1353–1365.

66. Chistiakov, D.A., A.A. Melnichenko, A.N. Orekhov, and Y.V. Bobryshev, Paraoxonase and atherosclerosis-related cardiovascular diseases. Biochimie, 2017. 132: p. 19–27.

67. Dubois, B., H. Hampel, H.H. Feldman, P. Scheltens, P. Aisen, S. Andrieu, H. Bakardjian, H. Benali, L. Bertram, K. Blennow, K. Broich, E. Cavedo, S. Crutch, J.F. Dartigues, C. Duyckaerts, S. Epelbaum, G.B. Frisoni, S. Gauthier, R. Genthon, A.A. Gouw, M.O. Habert, D.M. Holtzman, M. Kivipelto, S. Lista, J.L. Molinuevo, S.E. O’Bryant, G.D. Rabinovici, C. Rowe, S. Salloway, L.S. Schneider, R. Sperling, M. Teichmann, M.C. Carrillo, J. Cummings, C.R. Jack, Jr., G. Proceedings of the Meeting of the International Working, A.D. the American Alzheimer’s Association on “The Preclinical State of, July, and U.S.A. Washington Dc, *Preclinical Alzheimer’s disease: Definition, natural history, and diagnostic criteria*. Alzheimers Dement, 2016. 12(3): p. 292–323.

68. Cummings, J., P.S. Aisen, B. DuBois, L. Frolich, C.R. Jack, Jr., R.W. Jones, J.C. Morris, J. Raskin, S.A. Dowsett, and P. Scheltens, Drug development in Alzheimer’s disease: the path to 2025. Alzheimers Res Ther, 2016. 8: p. 39.

69. Dayon, L., A. Nunez Galindo, J. Wojcik, O. Cominetti, J. Corthesy, A. Oikonomidi, H. Henry, M. Kussmann, E. Migliavacca, I. Severin, G.L. Bowman, and J. Popp, Alzheimer disease pathology and the cerebrospinal fluid proteome. Alzheimers Res Ther, 2018. 10(1): p. 66.

70. Magistretti, P.J. and I. Allaman, A cellular perspective on brain energy metabolism and functional imaging. Neuron, 2015. 86(4): p. 883–901.

71. Malm, T., S. Loppi, and K.M. Kanninen, Exosomes in Alzheimer’s disease. Neurochem Int, 2016. 97: p. 193–9.

72. Wang, Y., V. Balaji, S. Kaniyappan, L. Kruger, S. Irsen, K. Tepper, R. Chandupatla, W. Maetzler, A. Schneider, E. Mandelkow, and E.M. Mandelkow, The release and trans-synaptic transmission of Tau via exosomes. Mol Neurodegener, 2017. 12(1): p. 5.

73. Saman, S., W. Kim, M. Raya, Y. Visnick, S. Miro, S. Saman, B. Jackson, A.C. McKee, V.E. Alvarez, N.C. Lee, and G.F. Hall, Exosome-associated tau is secreted in tauopathy models and is selectively phosphorylated in cerebrospinal fluid in early Alzheimer disease. J Biol Chem, 2012. 287(6): p. 3842–9.

74. Hong, S., V.F. Beja-Glasser, B.M. Nfonoyim, A. Frouin, S. Li, S. Ramakrishnan, K.M. Merry, Q. Shi, A. Rosenthal, B.A. Barres, C.A. Lemere, D.J. Selkoe, and B. Stevens, Complement and microglia mediate early synapse loss in Alzheimer mouse models. Science, 2016. 352(6286): p. 712–716.

75. Luchena, C., J. Zuazo-Ibarra, E. Alberdi, C. Matute, and E. Capetillo-Zarate, Contribution of Neurons and Glial Cells to Complement-Mediated Synapse Removal during Development, Aging and in Alzheimer’s Disease. Mediators Inflamm, 2018. 2018: p. 2530414.

76. Rajendran, L., M. Honsho, T.R. Zahn, P. Keller, K.D. Geiger, P. Verkade, and K. Simons, Alzheimer’s disease beta-amyloid peptides are released in association with exosomes. Proc Natl Acad Sci U S A, 2006. 103(30): p. 11172–7.

77. Bellingham, S.A., B.B. Guo, B.M. Coleman, and A.F. Hill, Exosomes: vehicles for the transfer of toxic proteins associated with neurodegenerative diseases? Front Physiol, 2012. 3: p. 124.

78. Dinkins, M.B., S. Dasgupta, G. Wang, G. Zhu, and E. Bieberich, Exosome reduction in vivo is associated with lower amyloid plaque load in the 5XFAD mouse model of Alzheimer’s disease. Neurobiol Aging, 2014. 35(8): p. 1792–800.

79. Montagne, A., Z. Zhao, and B.V. Zlokovic, Alzheimer’s disease: A matter of blood-brain barrier dysfunction? J Exp Med, 2017. 214(11): p. 3151–3169.

80. Shu, Y., R. Li, Y. Yang, Y. Dai, W. Qiu, Y. Chen, Z. Zhao, Z. Lu, and X. Hu, Urinary trypsin inhibitor levels are reduced in cerebrospinal fluid of multiple sclerosis and neuromyelitis optica patients during relapse. Neurochem Int, 2015. 81: p. 28–31.

81. Feng, M., Y. Shu, Y. Yang, X. Zheng, R. Li, Y. Wang, Y. Dai, W. Qiu, Z. Lu, and X. Hu, Ulinastatin attenuates experimental autoimmune encephalomyelitis by enhancing anti-inflammatory responses. Neurochem Int, 2014. 64: p. 64–72.

82. Mattsson, N., P.S. Insel, S. Palmqvist, E. Portelius, H. Zetterberg, M. Weiner, K. Blennow, O. Hansson, and I. Alzheimer’s Disease Neuroimaging, Cerebrospinal fluid tau, neurogranin, and neurofilament light in Alzheimer’s disease. EMBO Mol Med, 2016. 8(10): p. 1184–1196.

83. Mirra, S.S., A. Heyman, D. McKeel, S.M. Sumi, B.J. Crain, L.M. Brownlee, F.S. Vogel, J.P. Hughes, G. van Belle, and L. Berg, The Consortium to Establish a Registry for Alzheimer’s Disease (CERAD). Part II. Standardization of the neuropathologic assessment of Alzheimer’s disease. Neurology, 1991. 41(4): p. 479–86.

84. Braak, H. and E. Braak, Neuropathological stageing of Alzheimer-related changes. Acta Neuropathol, 1991. 82(4): p. 239–59.

85. Hyman, B.T. and J.Q. Trojanowski, Editorial on Consensus Recommendations for the Postmortem Diagnosis of Alzheimer Disease from the National Institute on Aging and the Reagan Institute Working Group on Diagnostic Criteria for the Neuropathological Assessment of Alzheimer Disease. Journal of Neuropathology & Experimental Neurology, 1997. 56(10): p. 1095–1097.

86. Brown, D.F., M.A. Dababo, E.H. Bigio, R.C. Risser, K.P. Eagan, C.L. Hladik, and C.L. White, III, Neuropathologic Evidence that the Lewy Body Variant of Alzheimer Disease Represents Coexistence of Alzheimer Disease and Idiopathic Parkinson Disease. Journal of Neuropathology & Experimental Neurology, 1998. 57(1): p. 39–46.

87. Boros, B.D., K.M. Greathouse, E.G. Gentry, K.A. Curtis, E.L. Birchall, M. Gearing, and J.H. Herskowitz, Dendritic spines provide cognitive resilience against Alzheimer’s disease. Annals of neurology, 2017. 82(4): p. 602–614.

88. Olsson, A., H. Vanderstichele, N. Andreasen, G. De Meyer, A. Wallin, B. Holmberg, L. Rosengren, E. Vanmechelen, and K. Blennow, Simultaneous Measurement of β-Amyloid (1–42), Total Tau, and Phosphorylated Tau (Thr181) in Cerebrospinal Fluid by the xMAP Technology. Clinical Chemistry, 2005. 51(2): p. 336–345.

89. Hulstaert, F., K. Blennow, A. Ivanoiu, H.C. Schoonderwaldt, M. Riemenschneider, P.P. De Deyn, C. Bancher, P. Cras, J. Wiltfang, P.D. Mehta, K. Iqbal, H. Pottel, E. Vanmechelen, and H. Vanderstichele, Improved discrimination of AD patients using beta-amyloid(1-42) and tau levels in CSF. Neurology, 1999. 52(8): p. 1555–62.

90. Shaw, L.M., H. Vanderstichele, M. Knapik-Czajka, C.M. Clark, P.S. Aisen, R.C. Petersen, K. Blennow, H. Soares, A. Simon, P. Lewczuk, R. Dean, E. Siemers, W. Potter, V.M. Lee, J.Q. Trojanowski, and I. Alzheimer’s Disease Neuroimaging, Cerebrospinal fluid biomarker signature in Alzheimer’s disease neuroimaging initiative subjects. Ann Neurol, 2009. 65(4): p. 403–13.

91. Mertins, P., L.C. Tang, K. Krug, D.J. Clark, M.A. Gritsenko, L. Chen, K.R. Clauser, T.R. Clauss, P. Shah, M.A. Gillette, V.A. Petyuk, S.N. Thomas, D.R. Mani, F. Mundt, R.J. Moore, Y. Hu, R. Zhao, M. Schnaubelt, H. Keshishian, M.E. Monroe, Z. Zhang, N.D. Udeshi, D. Mani, S.R. Davies, R.R. Townsend, D.W. Chan, R.D. Smith, H. Zhang, T. Liu, and S.A. Carr, Reproducible workflow for multiplexed deep-scale proteome and phosphoproteome analysis of tumor tissues by liquid chromatography-mass spectrometry. Nature protocols, 2018. 13(7): p. 1632–1661.

92. Yi, L., C.-F. Tsai, E. Dirice, A.C. Swensen, J. Chen, T. Shi, M.A. Gritsenko, R.K. Chu, P.D. Piehowski, R.D. Smith, K.D. Rodland, M.A. Atkinson, C.E. Mathews, R.N. Kulkarni, T. Liu, and W.-J. Qian, Boosting to Amplify Signal with Isobaric Labeling (BASIL) Strategy for Comprehensive Quantitative Phosphoproteomic Characterization of Small Populations of Cells. Analytical chemistry, 2019. 91(9): p. 5794–5801.

93. Russell, C.L., A. Heslegrave, V. Mitra, H. Zetterberg, J.M. Pocock, M.A. Ward, and I. Pike, Combined tissue and fluid proteomics with Tandem Mass Tags to identify low-abundance protein biomarkers of disease in peripheral body fluid: An Alzheimer’s Disease case study. Rapid Communications in Mass Spectrometry, 2017. 31(2): p. 153–159.

94. McKenzie, A.T., S. Moyon, M. Wang, I. Katsyv, W.-M. Song, X. Zhou, E.B. Dammer, D.M. Duong, J. Aaker, Y. Zhao, N. Beckmann, P. Wang, J. Zhu, J.J. Lah, N.T. Seyfried, A.I. Levey, P. Katsel, V. Haroutunian, E.E. Schadt, B. Popko, P. Casaccia, and B. Zhang, Multiscale network modeling of oligodendrocytes reveals molecular components of myelin dysregulation in Alzheimer’s disease. Molecular Neurodegeneration, 2017. 12(1): p. 82.

95. Tukey, J.W., Exploratory Data Analysis. 1977: Addison-Wesley.

96. Langfelder, P. and S. Horvath, Fast R Functions for Robust Correlations and Hierarchical Clustering. Journal of statistical software, 2012. 46(11): p. i11.

97. Yip, A.M. and S. Horvath, Gene network interconnectedness and the generalized topological overlap measure. BMC Bioinformatics, 2007. 8(1): p. 22.

98. Langfelder, P., B. Zhang, and S. Horvath, Defining clusters from a hierarchical cluster tree: the Dynamic Tree Cut package for R. Bioinformatics, 2008. 24(5): p. 719–20.

99. Miller, J.A., R.L. Woltjer, J.M. Goodenbour, S. Horvath, and D.H. Geschwind, Genes and pathways underlying regional and cell type changes in Alzheimer’s disease. Genome Med, 2013. 5(5): p. 48.

100. Zhang, Y., K. Chen, S.A. Sloan, M.L. Bennett, A.R. Scholze, S. O’Keeffe, H.P. Phatnani, P. Guarnieri, C. Caneda, N. Ruderisch, S. Deng, S.A. Liddelow, C. Zhang, R. Daneman, T. Maniatis, B.A. Barres, and J.Q. Wu, An RNA-sequencing transcriptome and splicing database of glia, neurons, and vascular cells of the cerebral cortex. J Neurosci, 2014. 34(36): p. 11929–47.

101. Sharma, K., S. Schmitt, C.G. Bergner, S. Tyanova, N. Kannaiyan, N. Manrique-Hoyos, K. Kongi, L. Cantuti, U.K. Hanisch, M.A. Philips, M.J. Rossner, M. Mann, and M. Simons, Cell type- and brain region-resolved mouse brain proteome. Nat Neurosci, 2015. 18(12): p. 1819–31.

